# CLUH maintains functional mitochondria and translation in motoneuronal axons and prevents peripheral neuropathy

**DOI:** 10.1101/2023.12.02.569712

**Authors:** Marta Zaninello, Tim Schlegel, Hendrik Nolte, Mujeeb Pirzada, Elisa Savino, Esther Barth, Hauke Wüstenberg, Tesmin Uddin, Lisa Wolff, Brunhilde Wirth, Helmar C. Lehmann, Jean-Michel Cioni, Thomas Langer, Elena I. Rugarli

## Abstract

Transport and local translation of mRNAs in distal axonal compartments are essential for neuronal viability. Local synthesis of nuclear-encoded mitochondrial proteins protects mitochondria from damage during their long journey along the axon, however the regulatory factors involved are largely unknown. Here, we show that CLUH, a cytosolic protein that binds mRNAs encoding mitochondrial proteins, is essential for preventing axonal degeneration of spinal motoneurons and maintaining motor behavior in the mouse. We demonstrate that CLUH is enriched in the growth cone of developing spinal motoneurons and is required for their growth. The absence of CLUH affects the abundance of target mRNAs and the corresponding mitochondrial proteins more prominently in axons, leading to ATP deficits specifically in the growth cone. CLUH binds ribosomal subunits, translation initiation and ribosome recycling components, and preserves axonal translation. Overexpression of the ribosome recycling factor ABCE1 rescues the growth cone and translation defects in CLUH-deficient motoneurons. In conclusion, we demonstrate a role for CLUH in mitochondrial quality control and translational regulation in axons, which are essential for their development and long-term integrity and function.

## Introduction

Neurons are highly polarized cells with distinct dendritic arborizations and axonal projections that can extend for long distances. Efficient trafficking of mitochondria and robust quality control mechanisms are essential to maintain the supply and the functionality of these organelles at distal sites, such as the growing tips of developing axons and the synaptic terminals in the mature nervous system (*1*). The long journey of mitochondria to reach peripheral positions makes them susceptible to depletion of short-lived proteins, less flexible to meet metabolic demands via proteome adaptation, and at risk of unbalanced stoichiometry of nuclear- and mitochondrially-encoded subunits of the respiratory chain (*1*). Local translation of mitochondrial proteins in axons is a safeguard mechanism to counteract mitochondrial ageing and preserve their function (*2–4*). mRNAs for mitochondrial proteins can be detected in axons, where they are associated with organelles, such as mitochondria and endosomes (*5–8*), or in RNA granules in complex with RNA-binding proteins (RBPs) (*9*). Several mRNAs for mitochondrial proteins were found to belong to a category of highly translated mRNAs in axons in the adult mammalian nervous system (*10*). However, knowledge of the ribonucleoprotein code that regulates the axonal transport, stability, and translation of these mRNAs is still sparse. Elucidating the involved molecular components is key to understand how neurons maintain a functional pool of mitochondria throughout a lifetime.

Clustered mitochondria homologue (CLUH) is an evolutionary conserved cytosolic RBP, which binds to a repertoire of mRNAs encoding mitochondrial proteins involved in oxidative phosphorylation (OXPHOS), the TCA cycle, and catabolic pathways (*11, 12*). In several organisms, CLUH loss induces mitochondrial clustering, reshapes the mitochondrial proteome, and leads to respiratory defects (*11, 13–18*). Constitutive lack of CLUH in human cells and mouse liver is associated with a downregulation of its target mRNAs, which are subjected to enhanced decay (*16, 19*). *In vivo*, CLUH function is crucial at metabolic switches, such as the fetal-neonatal transition, starvation, or adipogenesis (*16, 20*). Although several lines of evidence suggest a role for CLUH in translational regulation (11, 18, 20, 21), how this occurs is an open question. Here, we examined the role of CLUH in neurons whose integrity depends on RNA transport and local translation. Ablation of *Cluh* expression in mouse neural progenitors revealed a role in maintaining motor behavior, by preventing late-onset degeneration of distal axons and neuromuscular junctions (NMJs). *In vitro*, CLUH is essential for axonal growth, maintains functional mitochondria at the growth cone (GC), and sustains axonal translation. CLUH associates with components involved in initiation of translation and ribosome recycling and is essential to preserve these components in axons. We find that overexpression of ABCE1, which is involved in ribosomal quality control and translational initiation, rescues the growth cone size and the axonal translational defect observed in CLUH-deficient motoneurons. Our data link CLUH to translational quality control and demonstrate the importance of this process in maintaining functional mitochondria in distal axons.

## Results

### *Cluh* deletion in the mouse neural progenitors impairs locomotor activity and causes peripheral neuropathy

Cortical and spinal motoneurons, which control voluntary movements, are endowed with the longest axons in the central nervous system and are prone to degenerate when local translation or mitochondrial function is impaired (*1, 21, 22*). To investigate if CLUH is required for the maintenance of neuronal circuits *in vivo,* we knocked out the gene in mouse neural progenitors, by crossing previously characterized mice carrying *Cluh* alleles with a loxP-flanked exon 10 (*16*) with nestin-Cre transgenic mice (*23*) (genotype: *Cluh*^fl/fl^ nestin-Cre^tg/+^ from now on indicated as NKO). CLUH expression was successfully ablated in the forebrain and cerebellum and substantially reduced in the spinal cord (Fig. S1A). NKO mice were viable, fertile, but had a smaller body weight, and a slightly increased brain weight when normalized by the body weight (Fig. 1A-C). Motor behavior was normal at 4 months, but at 5 months NKO mice showed locomotor impairment, as revealed by a classical walking beam test, a proxy of the motor coordination of hindlimbs (Fig. 1D-G and Fig. S1B, C). NKO mice were slower, and slipped more frequently while crossing the beam, independently of their gender (Fig. 1D-G, and Supplementary Videos 1-3). The effect of *Cluh* deletion was more severe 3 months later (Fig. 1D-G, and Supplementary Videos 1-3), indicating a progressive phenotype. The motor deficit was confirmed by a reduced latency time of NKO mice to fall in the accelerating rotarod test, which measures grip, motor coordination and balance, with males showing a more severe phenotype than females in this test (Fig. S1B, C). In all experiments, nestin-Cre mice (Cre) were also tested to exclude any effect due to the expression of the Cre recombinase (*24*).

**Fig. 1:**
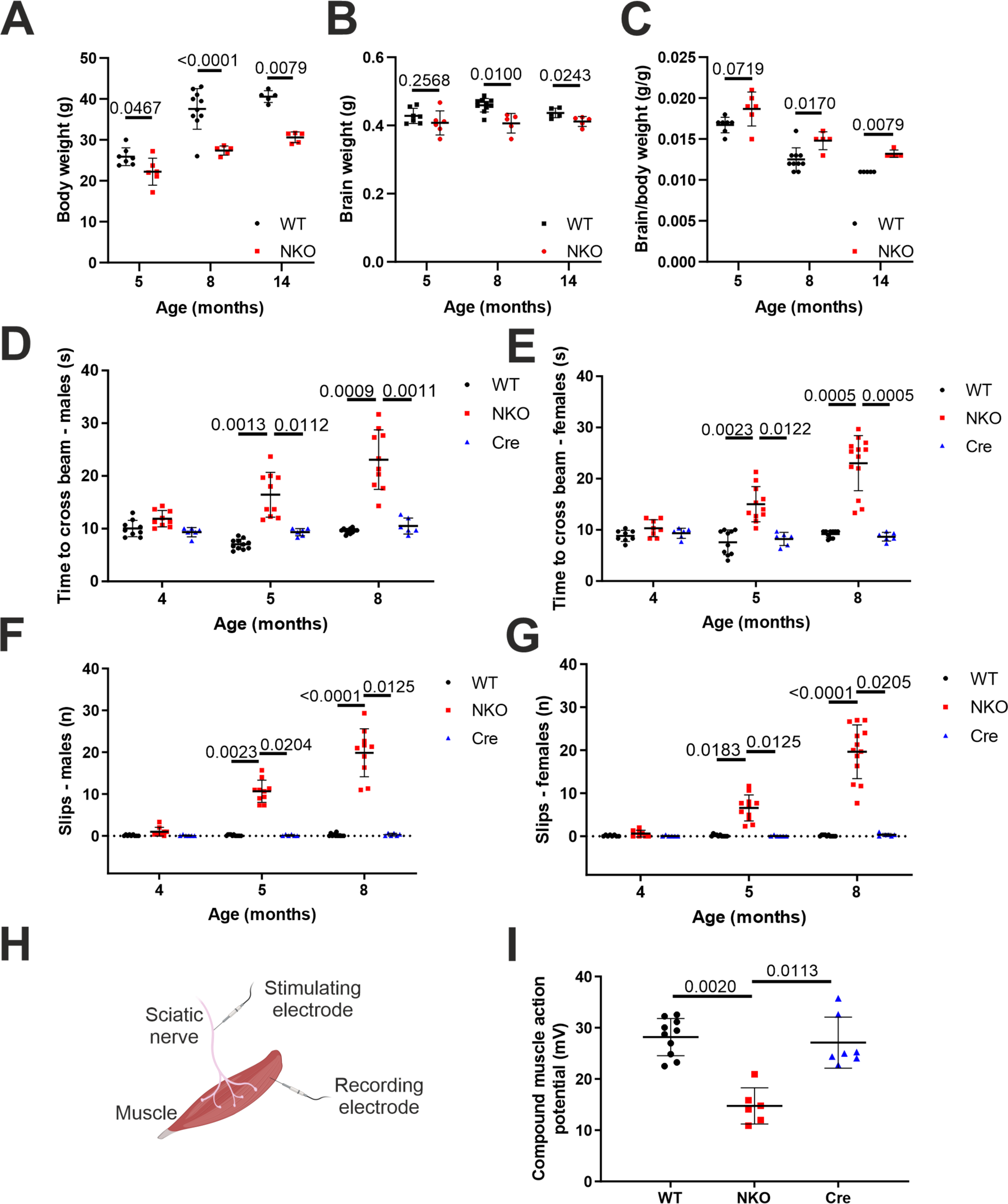
NKO mice show locomotor defects (**A-C**) Body weight (A), brain weight (B) and brain/body weight ratio (C) of WT and NKO mice at 5, 8 and 14 months of age. Data represent mean ± SD of 5-10 mice. Statistical significance was determined by by Mann-Whitney and Welch’s t tests. (**D-G**) Quantification of the walking beam test in males (D, F) and females (E, G) aged 4, 5 and 8 months as time spent to cross the beam (D, E) and number of total slips during the crossing (F, G). Data represent mean ± SD of 6-13 mice. Statistical significance was determined by by Mann-Whitney and Welch’s t tests. (**H**) Scheme depicting CMAP recording. Sciatic nerve was stimulated with an electrode and the action potential was measured with a recording electrode in the muscle of the hind paw. (**I**) Quantification of the CMAP amplitude recorded in the hind paw after stimulation of the sciatic nerve of male mice aged 8 months. Data represent mean ± SD of 6-10 mice. Statistical significance was determined by Dunn’s multiple comparison test.

The motor impairment of NKO mice suggested a peripheral neuropathy. To test this hypothesis, we performed electrophysiological studies to record the sum of actions potentials (compound muscle action potential, CMAP) generated in the muscles of the hind paw, after stimulating the sciatic nerve that contains the axons of spinal motoneurons. The CMAP amplitude, which is a reliable index of axonal damage, was halved in NKO muscles compared to WT muscles at 8 months (Fig. 1H, I). To substantiate these findings, we collected the distal part of the sciatic nerve, where it separates in the tibial and peroneal branches and performed semithin sections at earlier and later ages. While at 1 month of age the NKO nerves contained a similar number of axons of comparable size (Fig. S2A-D), at 5 and 14 months we observed a pathological phenotype especially pronounced in the peroneal branch of the NKO nerve. This was characterized by a decreased number of axons per area, with an increase of smaller axons and a decrease of large axons, and the appearance of profiles suggesting axonal degeneration (Fig. 2A- D). The pathological phenotype was very similar at 14 months of age (Fig. 2A-D).

**Fig. 2:**
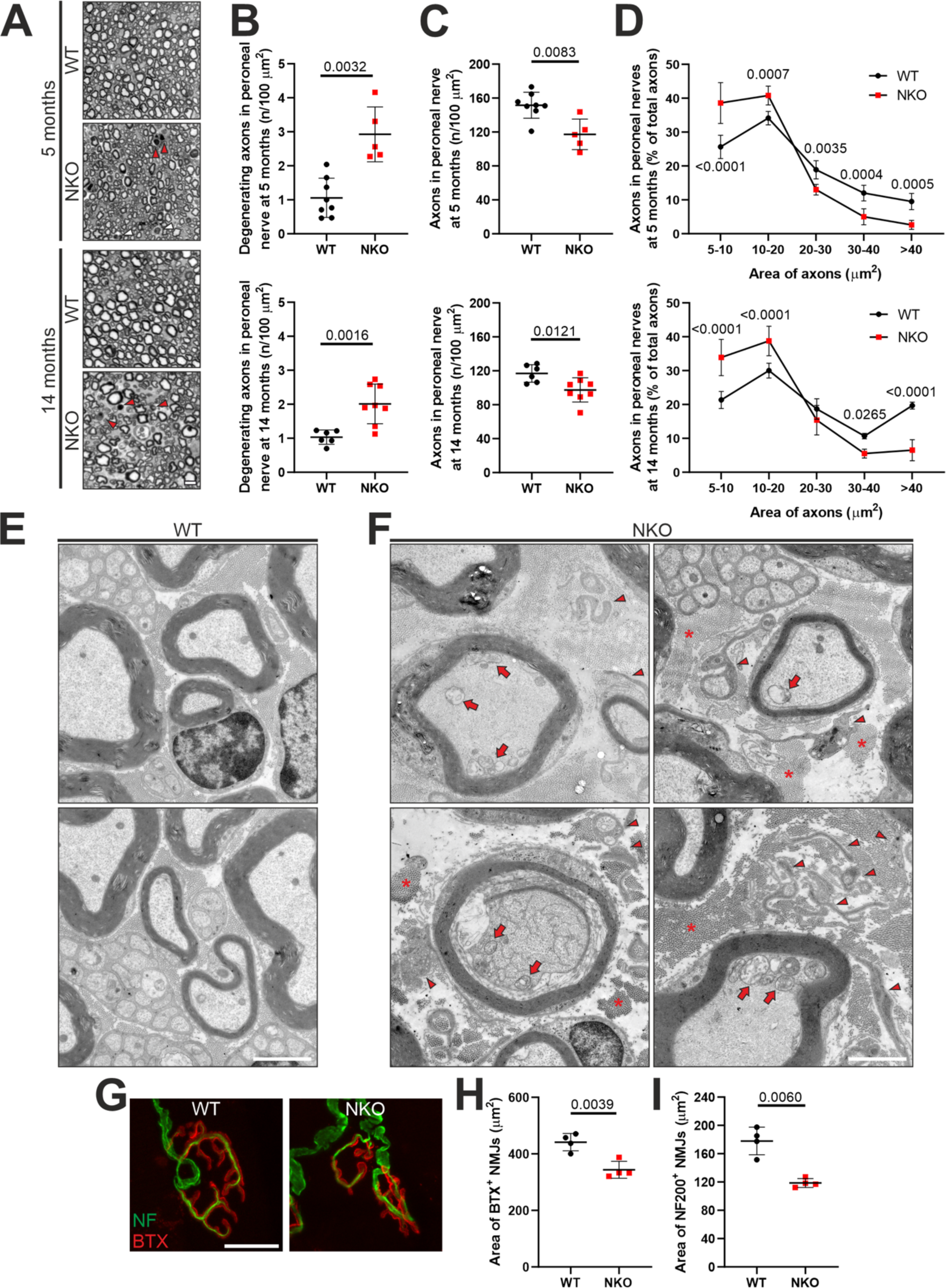
NKO mice show peripheral neuropathy (**A**) Semithin sections of the peroneal branch of the sciatic nerve of 5 and 14 months-old mice. Arrowheads indicate degenerating axons. Bar, 5 µm. (**B-D**) Quantification of degenerating axons (B), of the number of axons larger than 5 µm^2^ per area (C), and of the distribution of axons of different diameters (D) in semithin sections of the peroneal branch of the sciatic nerve at indicated ages. Data represent mean ± SD of 5-8 mice. Statistical significance was determined by Welch’s t test in B, C and by Sidak’s multiple comparisons test in D. (**E, F**) Electron micrographs of peroneal nerves of WT (E) and NKO (F) mice aged 14 months. Arrows indicate abnormal swollen mitochondria, asterisks show accumulation of collagen, and arrowheads point to denervated Schwann cells. Bar, 2 µm. (**G**) NMJs of transverse abdominal muscles in WT and NKO mice aged 14 months. NF, Neurofilament; BTX, α-bungarotoxin. Bar, 25 µm. (**H, I**) Quantification of the BTX^+^ area of post synapses (H) and NF^+^ area of pre synapses (I) in experiments as in (G). Data represent the mean ± SD of 4 mice (43-91 NMJs per mouse). Statistical significance was determined by Welch’s t test.

Ultrastructurally, the NKO nerves were characterized by basal lamina–covered Schwann cell profiles devoid of axons and myelin that most probably represent denervated Schwann cells (reminiscent of bands of Büngner), increased connective tissue with collagen deposition, and collagen pockets, which are more often observed with loss of small unmyelinated fibers (Fig. 2E, F). Moreover, we found myelinated fibers with accumulated organelles or containing swollen mitochondria, indicating impaired axonal trafficking preceding degeneration (Fig. 2F).

Mitochondria in NKO axons showed more frequently open cristae than in WT (Fig. S2E, F). Axonal pathology was supported by transcriptome changes detected in the sciatic nerve at 5 months of age. Transcripts involved in neurotransmitter regulation and synaptic function were decreased in NKO nerves, whereas mRNAs coding for pro-survival and inflammatory components were upregulated, in agreement with a degenerative program (Table S1, Fig. S2G).

To further characterize the phenotype of motor axons, we analyzed the NMJs, specialized synapses of spinal motoneurons with the skeletal muscle, whose size affects motor performance and diminishes with aging and motor neuron diseases (*25*). The innervation and postsynaptic areas of NKO NMJs were smaller than in WT mice at 14 months of age (Fig. 2G-I). In contrast, the number of spinal motoneurons positive for choline acetyltransferase (ChAT) was unaffected in NKO spinal cords (Fig. S2H, I), indicating that loss of CLUH induces specifically an axonopathy. Taken together, our data identify a crucial role of CLUH *in vivo* to support motor behavior, preserve motor axon function in the sciatic nerve, and prevent loss of axons.

### CLUH regulates the size and the mitochondrial ATP levels of growth cones of spinal motoneurons

To understand if the axonal phenotype triggered by loss of CLUH *in vivo* is cell-autonomous, we turned to *in vitro* experiments using primary embryonic spinal motoneurons cultured in absence of glia. After 6 days *in vitro* (DIV), NKO motoneurons displayed shorter axons compared to WT motoneurons (Fig. 3A, B), and at DIV 10 started to die, as assessed by a decreased number of TAU^+^ and increased number of TUNEL^+^ neurons (Fig. S3A-D). For this reason, we performed all subsequent experiments in vitro at DIV 6. CLUH levels did not change during neuronal maturation in *vitro* (Fig. S3E) and the expression of the motoneuronal markers Hb9 and ChAT (*26*) was not affected in absence of CLUH (Fig. S3F). Neither the axonal density of mitochondria, nor the neuronal mt-DNA levels were affected by loss of CLUH (Fig. S4A-C). Moreover, WT and NKO mitochondria in axons were comparable for morphology and membrane potential (*27*) (Fig. S4D, E). However, the size of GCs and their content of synaptic vesicles was reduced in NKO motoneurons (Fig. 3C-F), consistent with a growth defect. To control for the specificity of this phenotype, we transfected primary motoneurons with a vector expressing the mouse isoform of CLUH tagged with a mCherry fluorescent moiety at the C-terminus (Fig. S5A). Overexpressed CLUH was highly enriched in the soma, but was also present along the axons, especially close to axonal protrusions, and was enriched in the peripheral part of GCs, which contains tyrosinated tubulin, IQGAP1 and CLASP2, but not acetylated tubulin (*28*) (Fig. S5B, C). We confirmed that an untagged human CLUH construct was similarly located at the GC (Fig. S5D), and that an empty mCherry vector did not produce the same GC staining (Fig. S5E). Importantly, overexpression of CLUH rescued both axonal length and GC size of NKO motoneurons (Fig. 3G- I). The axonal grow phenotype observed *in vitro* prompted us to evaluate if axonal growth defects occurred also *in vivo* as a delay in innervation, which may be later compensated. Notably, developing axons in the hind limbs labeled by neurofilament antibodies in whole-mount E13.5 NKO embryos were shorter than in WT (Fig. 3J, K).

**Fig. 3:**
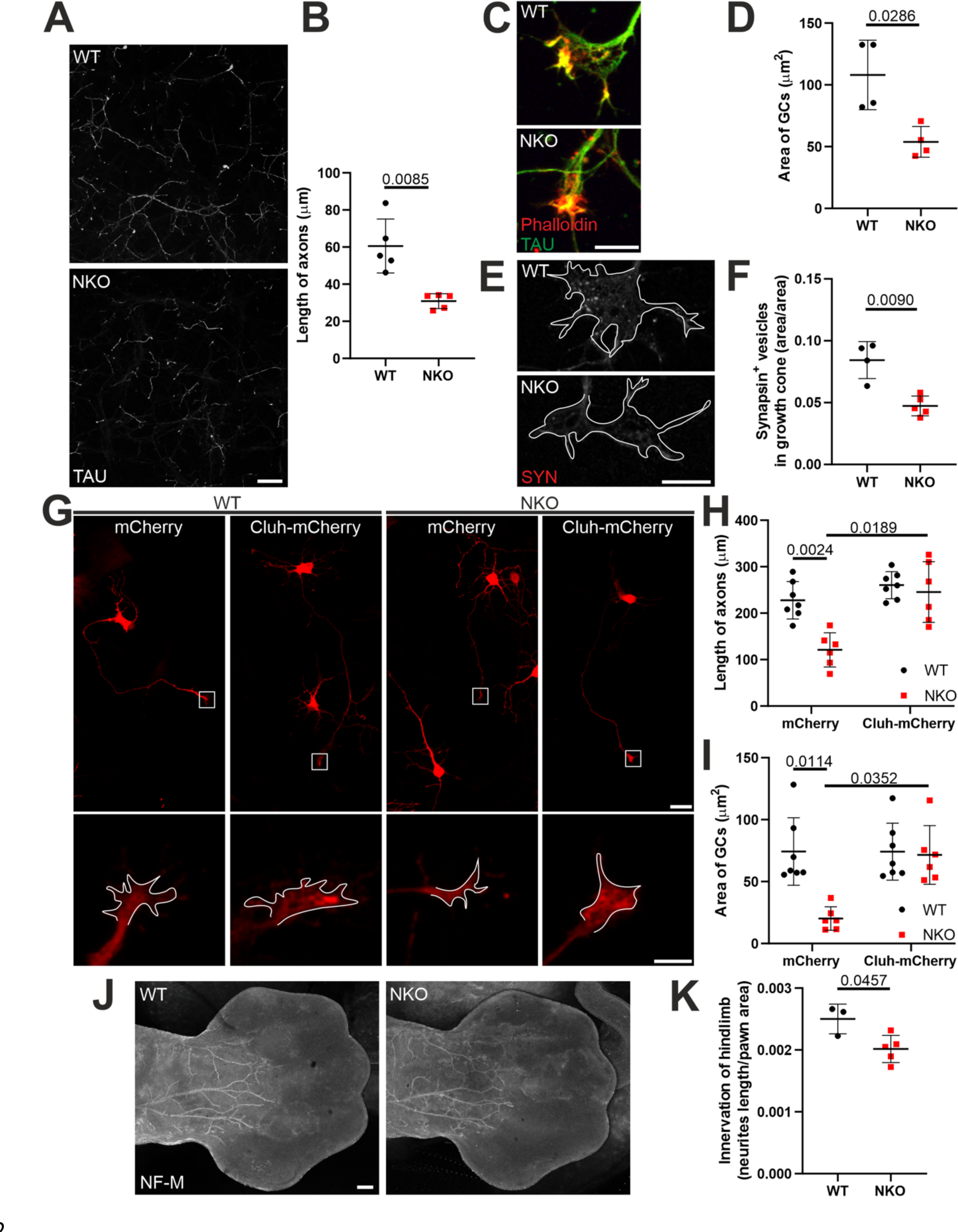
Lack of CLUH impairs axonal growth (**A**) Motoneuronal axons stained by TAU in the axonal compartment of Boyden chambers at DIV 6. Bar, 75 µm. (**B**) Quantification of the length of axons in experiments as in (A). Data represent the mean ± SD of 5 mice (4-5 fields per mouse). Statistical significance was determined by Welch’s t test. (**C**) GCs of primary spinal motoneurons stained for TAU (green) and actin (Phalloidin, red). Bar, 10 µm. (**D**) Quantification of the area of GCs in experiments as in (C). Data represent the mean ± SD of 4 mice (47-78 GCs per mouse). Statistical significance was determined by Mann-Whitney test. (**E**) Staining of SYNAPSIN^+^ (SYN) vesicles in GCs of primary spinal motoneurons. GCs are outlined in white. Bar, 10 µm. (**F**) Quantification of the number of SYN^+^ vesicles in GCs in experiments as in (E). Data represent the mean ± SD of 4-5 mice (26-35 GCs per mouse). Statistical significance was determined by Welch’s t test. (**G**) WT and NKO primary motoneurons expressing mCherry or Cluh-mCherry. GCs are magnified in the bottom panels. Bars, 20 µm and 5 µm. (**H, I**) Quantification of the length of axons (H) and size of GCs (I) in experiments as in (G). Data represent the mean ± SD of 6-7 mice (6-14 axons or GCs per mouse). Statistical significance was determined by Dunn’s and Dunnett’s multiple comparison tests. (**J**) Whole mounts of hindlimbs of E13.5 WT and NKO embryos stained with neurofilament-M (NF-M). Bar, 200 µm. (**K**) Quantification of the innervation of the hind limbs in respect to the area of the paw as in (J). Embryos belonged to the same litter. Data represent the mean ± SD of 3-5 mice. Statistical significance was determined by Welch’s t test.

GCs dynamically remodel the cytoskeleton in response to attractive and repulsive chemical guidance clues. This process is supported by cycles of de- and re-polymerization of actin and microtubules which require boosts of energy. Since CLUH deficiency impairs mitochondrial respiration (*16, 17, 19*), insufficient ATP production from mitochondria may underlie the distal defects of NKO motoneurons. We measured mitochondrial ATP in soma and GCs using the ratiometric sensor ATeam targeted to mitochondria (*29*). ATP levels were decreased in mitochondria of NKO GCs compared to WT but were unaffected in soma mitochondria (Fig. 4A- C). Blockage of the ATP synthase by oligomycin reduced the levels of mitochondrial ATP in WT but decreased it only slightly in NKO GCs (Fig. 4C). Cytosolic ATP levels were also reduced in NKO GCs when neurons were cultured in galactose medium to limit glycolysis rate (Fig. 4D).

**Fig. 4:**
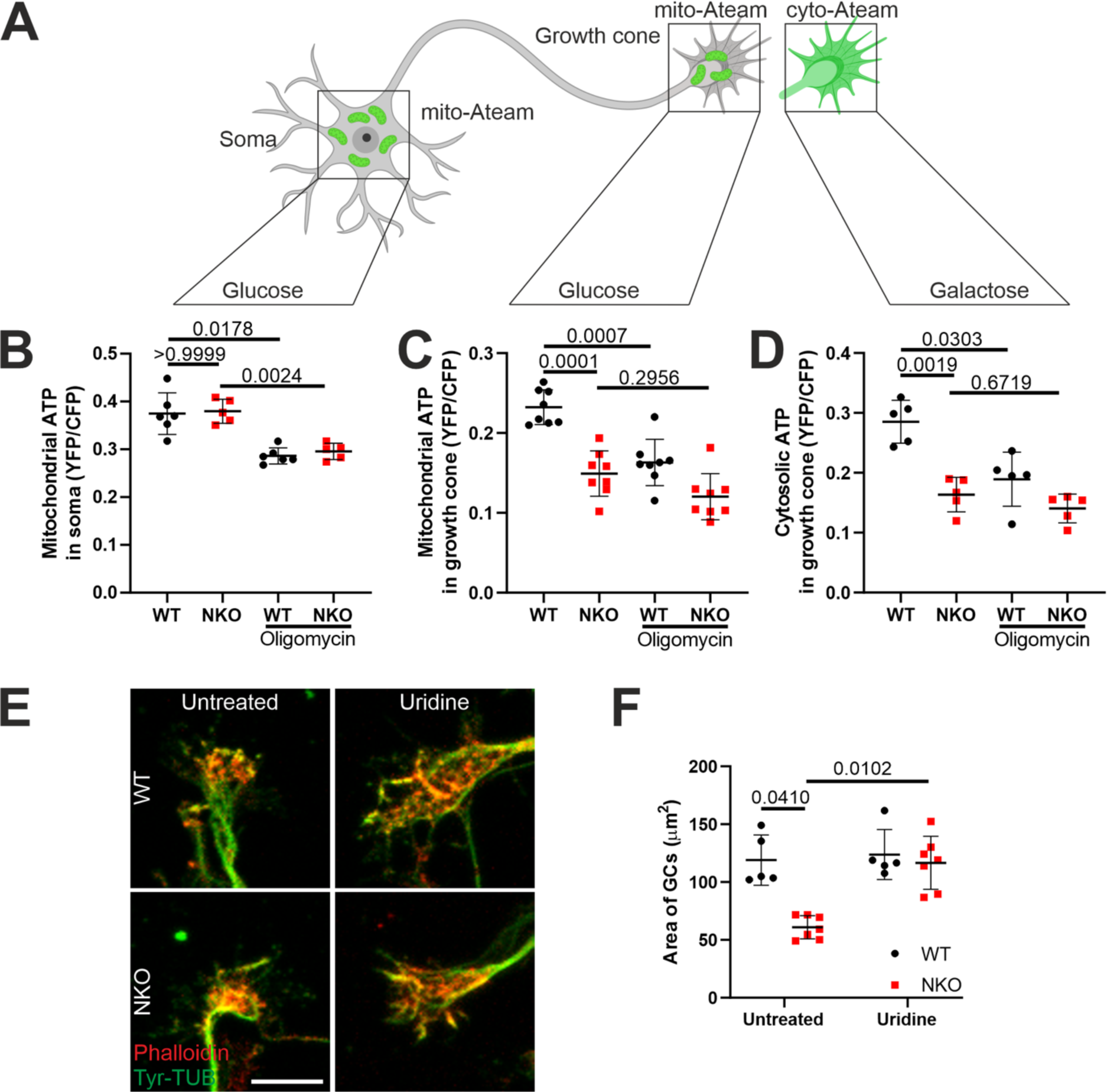
Lack of CLUH leads to ATP defects in GCs (**A**) Depiction of the experimental strategy using the ATP sensor mito-ATeam targeted to mitochondria or the cytosol. (**B, C**) Quantification of mitochondrial ATP in somas (B) and GCs (C) of primary motoneurons transfected with mito-ATeam. The analyzed somas are from the same field of GCs. Data represent the mean ± SD of GCs and somas from 5-8 mice. Statistical significance was determined by Dunnett’s multiple comparison test. (**D**) Quantification of cytosolic ATP in GCs of primary motoneurons transfected with the cytosolic ATeam sensor and grown in galactose. Data represent the mean ± SD of GCs from 5 mice. Statistical significance was determined by Dunnett’s multiple comparison test. (**E**) GCs stained with Tyr-TUB (green) and Actin (Phalloidin, red) of motoneurons grown in medium with or without uridine supplementation. Bar, 10 µm. (**F**) Quantification of the size of GCs in experiments as in (E). Data represent the mean ±SD of 5-7 independent cultures (27-66 GCs per culture). Statistical significance was determined by Dunn’s multiple comparison test.

Moreover, the acute inhibition of mitochondrial ATP synthase by oligomycin induced the collapse of WT GCs to a greater extent than NKO GCs (Fig. S6A, B). Thus, CLUH maintains mitochondrial ATP levels in GCs.

Glycolysis metabolically supports distal axons and growth cone dynamics (*30, 31*), raising the question whether the impaired mitochondrial ATP production observed in NKO GCs is at the root of the phenotype. Loss of CLUH broadly affects mitochondrial metabolism, including nucleotide synthesis (*17, 19*). The mitochondrial rate-limiting enzyme in pyrimidine synthesis, DHODH, uses coenzyme Q as electron acceptor and is therefore dependent on a functional respiratory chain (*32*). For this reason, respiratory deficient cells cannot grow without uridine supplementation (*33, 34*). Uridine is a precursor of RNA and phospholipids and can be converted in ribose-1-phosphate that enter glycolysis to fuel ATP production (*35*). We therefore tested if boosting ATP levels by uridine supplementation could recover the GC phenotype in NKO axons. Notably, the area of NKO GCs increased in presence of uridine (Fig. 4E, F), supporting the hypothesis that a metabolic deficit of distal mitochondria is one of the causes of collapsed GCs.

### CLUH controls the abundance of target mRNAs in axons

CLUH may affect mitochondrial respiration and metabolism by regulating target mRNAs at different levels. We first checked if the abundance of CLUH target mRNA molecules was similarly affected in cell bodies and axons of primary motoneurons, and selected *Atp5a1* and *Pink1* for further analysis by FISH. *Atp5a1* mRNA molecules were reduced in both NKO cell bodies and axons, but this reduction was more pronounced in axons, in agreement with the distal ATP defect (Fig. 5A-D). Consistenly, *Pink1* mRNA molecules were also decreased in the axon in absence of CLUH. *ActB* mRNA, which is not bound by CLUH, was not altered in NKO somas and axons (Fig. 5A-D). Notably, the distribution of mRNA molecules for all transcripts analyzed was similar in WT and NKO axons (Fig. 5E-G).

**Fig. 5:**
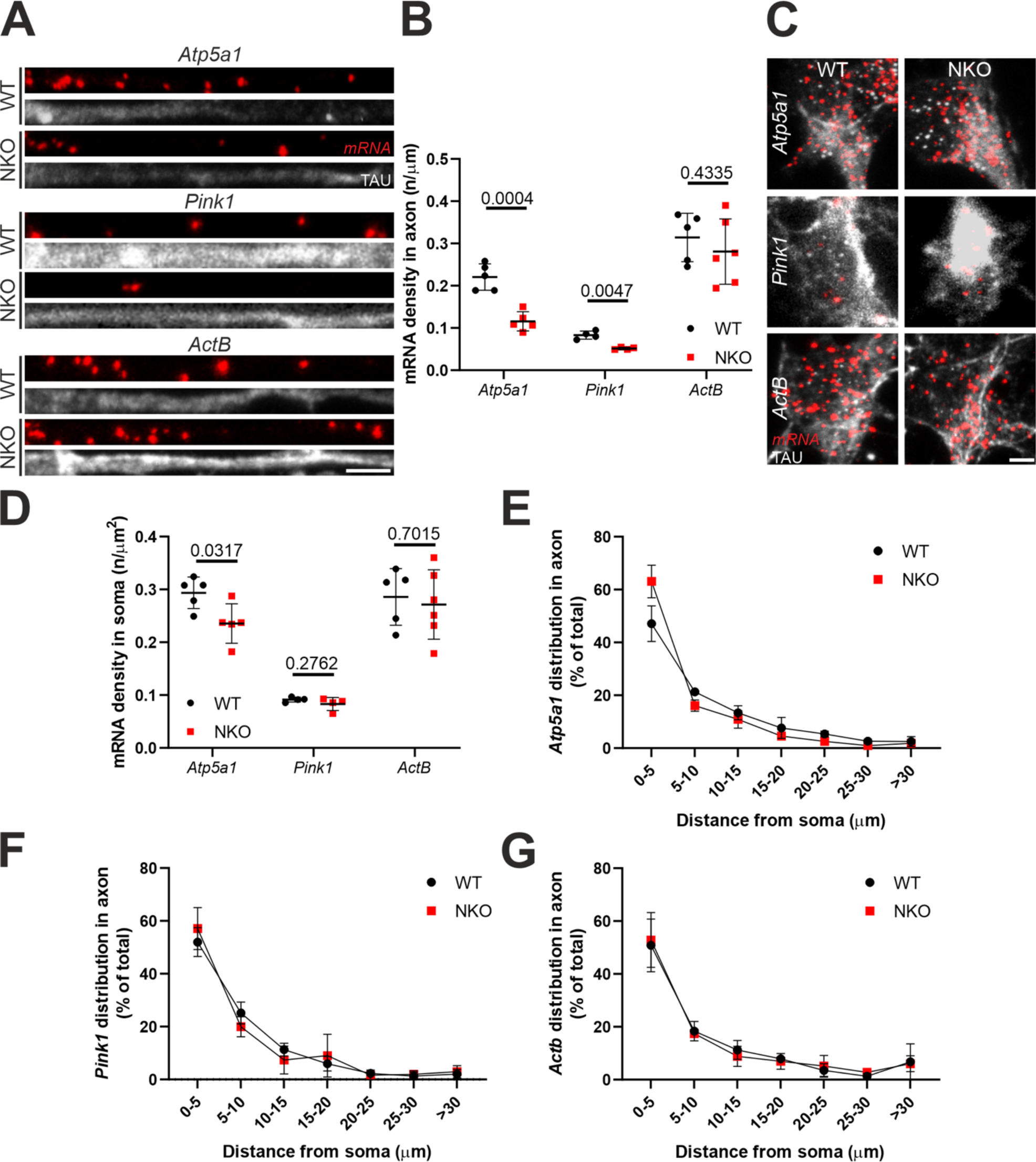
Lack of CLUH depletes axons and somas of CLUH target mRNAs (**A**) RNAscope of *Atp5a1*, *Pink1* and *ActB* mRNAs (red) in axons of primary spinal motoneurons (DIV 6) stained for TAU (grey). Bar, 5 µm. (**B**) Quantification of the abundance of *Atp5a1*, *Pink1* and *ActB* mRNAs in axons in experiments as in (A). Data represent the mean ± SD of 4-6 mice (26-57 neurons per mouse). Statistical significance was determined by Welch’s t test. (**C**) RNAscope of *Atp5a1*, *Pink1* and *ActB* mRNAs (red) in axons of primary spinal motoneurons (DIV 6) stained for TAU (grey). Bar, 5 µm. (**D**) Quantification of the abundance of *Atp5a1*, *Pink1* and *ActB* mRNAs in somas in experiments as in (C). Data represent the mean ± SD of 4-6 mice (26-57 neurons per mouse). Statistical significance was determined by Mann-Whitney and Welch’s t test. (**E**-**G**) Distribution of *Atp5a1* (E), *Pink1* (F) and *ActB* (G) mRNAs particles along axons of primary motoneurons in experiments as shown in (A). Data represent the mean ± SD of 3-5 cultures (72-653 mRNA dots per culture).

One possible explanation for the previous findings is that mRNAs are less transported into axons when not bound by CLUH. To directly visualize trafficking of mRNAs in WT and NKO motoneurons, we used the bacteriophage derived MS2 tether system. The CLUH mRNA targets *Atp5a1* and *Mdh2*, and the *Actb* mRNA as a control were tagged after the 3’ UTR with 24 repeats of the MS2 stem-loops and expressed in neurons together with a reporter construct encoding the MS2 coat protein (MCP) fused with a Halo tag for fluorescent detection (Fig. S7A). For analysis, neurons showing axonal mRNA particles were selected. The time in motion of *Atp5a1*, *Mdh2*, and *Actb* tagged mRNAs was similar in WT and NKO axons (Fig. S7B, C). Furthermore, we did not find significant changes in the type of movement of the mRNA particles, assessed either by lateral displacement over the whole track (directed, oscillatory, or stationary) (Fig. S7D) or by the mean square displacement in the first 25% of the track (active, diffusive, or confined) (Fig. S7E). Thus, the reduced axonal localization of CLUH target mRNAs is likely caused by a decrease of mRNA stability rather than a trafficking defect along the axons.

### Compartmentalized proteomics reveals mitochondrial and translational signatures in NKO axons

To further unravel the underlying cause for the axonal phenotype, we examined whether CLUH depletion affects the proteome differently depending on the neuronal compartments. To this end, we isolated the central and the peripheral portion of neurons using modified Boyden chambers (*36*) and performed proteomics analysis (Fig. 6A). Motoneurons were grown on the porous membrane in the upper compartment of the chamber (from now on named neuron compartment) and only neurites could pass into the bottom compartment. Since embryonic motoneurons have very short dendrites, the bottom compartment is highly enriched of axons (from now on named axon compartment, Fig. S8A). Primary motoneurons were isolated from different embryos in independent experiments and grown in Boyden chambers for six days before collecting the two compartments. Given the paucity of material of the axonal compartment, two samples of the same genotype and neuronal culture were pooled before mass spectrometry (MS) analysis. The proteomics of the axonal compartment relative to the neuron compartment revealed an enrichment of axonal markers and a depletion of proteins located in the nucleus, ER and Golgi (Fig. S8B), confirming the quality of the separation. Since principal component analysis showed a batch effect associated with the day of neuronal isolation and the litter (Fig. S8C, D), data were transformed to log2 fold changes of NKO versus WT samples for each individual litter and neuronal isolation followed by a one-sample t-test. Proteins were included in the analysis if quantified in at least 3 out of 7 NKO-WT comparisons. Accordingly, 2,800 proteins were identified in the axonal and 6,453 in the whole neuronal compartment (Supplementary Table 2). In the NKO neuron-containing fractions, 52 proteins were increased (log2FC > 0.4; p-value < 0.05) and 117 were decreased (log2FC < -0.4; p-value < 0.05) relative to WT. In the NKO axonal fractions, 14 proteins were upregulated (log2FC > 0.4) and 60 proteins were downregulated (log2FC < -0.4) (Fig. 6B). In both compartments, GO analysis did not identify any specific upregulated pathway, but showed an over-representation of mitochondrially-related terms among the downregulated proteins (Fig. 6C, D, Fig. S8E). Thus, mitochondrial proteins were decreased centrally and distally in the absence of CLUH. We observed a significant overlap between mitochondrial proteins whose steady state level was decreased and those encoded by CLUH mRNA targets (*11*) (Fig. S8F). Of note, these were more decreased in axons with respect to the whole neurons (Fig. 6E). These proteins included components of the respiratory chain, TCA cycle, β-oxidation and other metabolic enzymes.

**Fig. 6:**
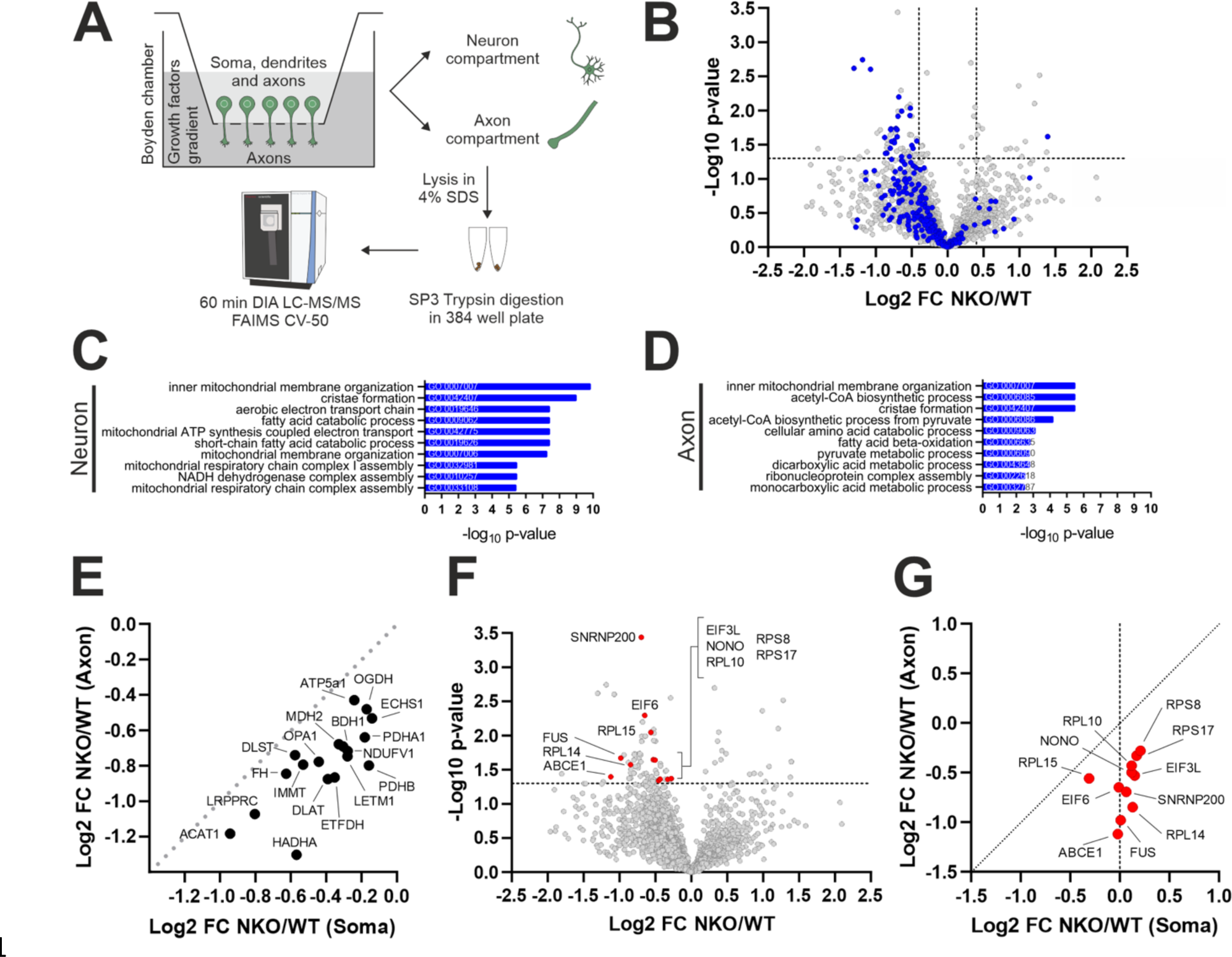
Mitochondrial proteins and translation components are depleted in NKO axons (**A**) Experimental setup to isolate axons of spinal motoneurons using Boyden chambers. A gradient of growth factors was applied from the lower axonal compartment to the upper compartment containing soma, dendrites, and axons. Axons were scraped and lysed in a 4% SDS solution; immediately afterwards the upper compartment was lysed in the same solution. Samples were then digested in SP3 Trypsin buffer and subsequently analyzed using DIA LC-MS/MS FAIMS CV-50. (**B**) Volcano plot of protein changes in NKO versus WT axons. Mitochondrial proteins are highlighted in blue. (**C, D**) GOBP analysis of downregulated proteins in the neuron (C) and axon (D) compartments of primary motoneurons grown in Boyden chambers. Analysis was done using the EnrichR webtool. (**E**) Correlation of the fold change of proteins encoded by CLUH mRNA targets in NKO versus WT axon and neuron compartments. Only proteins measured in axons are indicated. (**F**) Volcano plot of proteins altered in NKO versus WT axons. Proteins related to translation or RNA binding are highlighted in red. (**G**) Correlation of the fold changes of proteins highlighted in (F) in NKO versus WT in axon and neuron compartments.

Besides mitochondrial proteins, proteins reduced in NKO axons included other RBPs (FUS, NONO), the splicing factor SNRNP200, crucial components for the initiation and regulation of translation (ABCE1, EIF3L, EIF6), and several ribosomal subunits (RPL10, RPL14, RPL15, RPS8, RPS15) (Fig. 6F). We confirmed these results by measuring the decreased intensity of the fluorescence of ABCE1, RPL14 and RPS8 in NKO-axons (Fig. S9A, B). In contrast to mitochondrial proteins, these translational components were unchanged in the compartment containing the cell bodies (Fig. 6G). Thus, while the mitochondrial proteomic signature is common to neuronal soma and axons, albeit more pronounced in axons, the translational signature was restricted to axons.

### Impaired axonal translation induced by loss of CLUH is rescued by ABCE1 overexpression

The previous findings prompted us to examine whether CLUH is required to preserve axonal translation. To this end, we employed fluorescent noncanonical amino acid tagging (FUNCAT), which uses the incorporation of the methionine analog homopropargylglycine (HPG) in nascent proteins and the subsequent labeling via click reaction with a fluorescent azide. Indeed, these experiments established that defects of translational components were reflected in reduced protein synthesis in NKO axons (Fig. 7A, B), but not in somas (Fig. 7C, D). It is known that mitochondrial proteins are highly translated in axons (*10*), however it is still surprising that lack of CLUH leads to a general defect of translation in axons. This result prompted us to investigate the underlying molecular mechanisms. We therefore immunoprecipitated endogenous CLUH in HeLa cells followed by MS to reveal its interactome. We found that several ribosomal components and translational regulators, including those depleted in NKO axons, were significantly enriched in the CLUH immunoprecipitation (IP) versus the control IP, demonstrating a link between CLUH and the translational machinery (Fig. 7E, F and Supplementary Table 3).

**Fig. 7:**
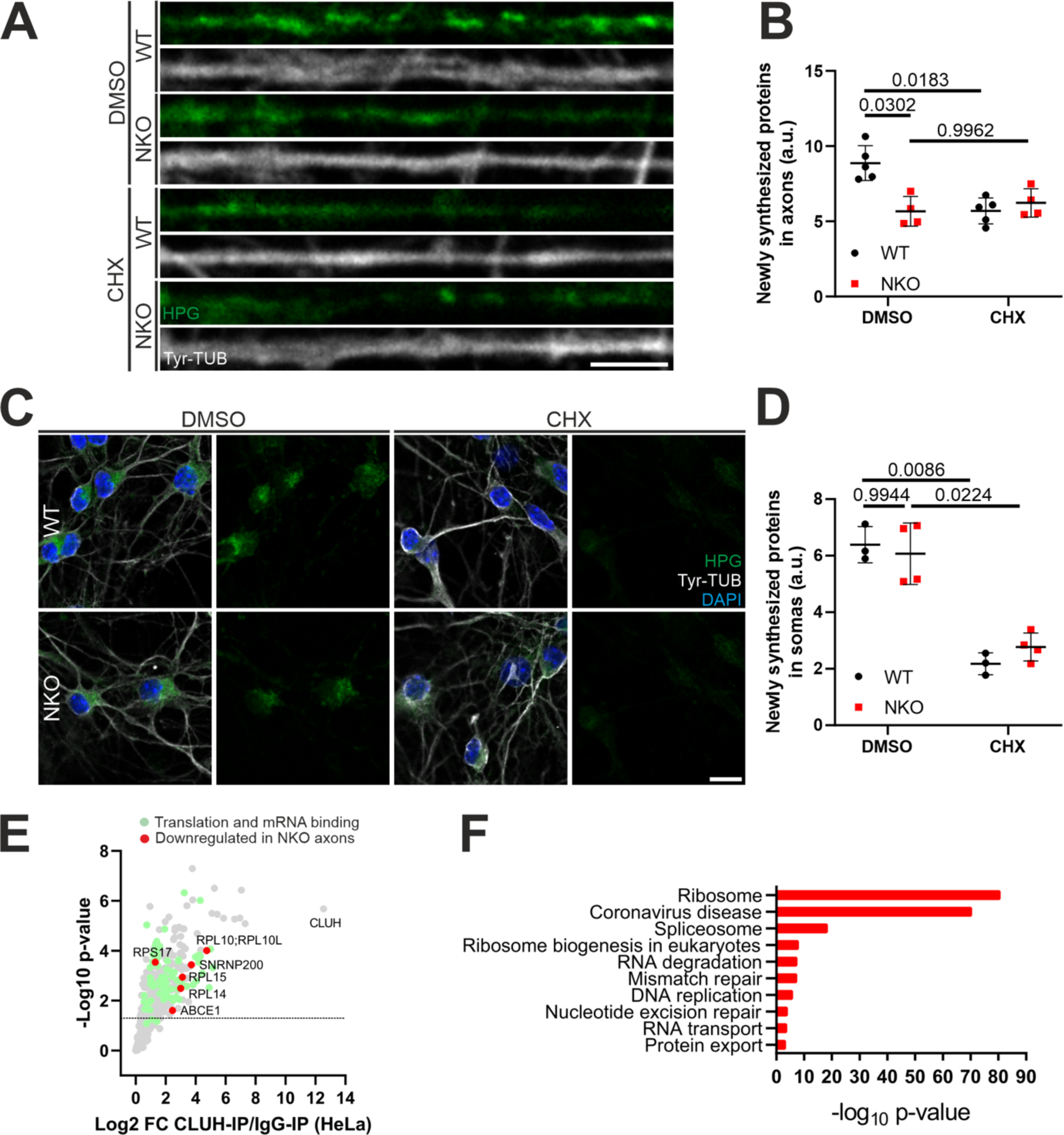
CLUH absence impairs translation in axons but not in somas (**A, B**) Single confocal planes (A) and quantification (B) of newly synthesized proteins (HPG, green) revealed by the FUNCAT assay in WT and NKO axons stained with Tyr-TUB (grey). Data represent the mean ± SD of 4-5 mice (27-71 axons per culture). Statistical significance was determined by Dunnett’s multiple comparison test. CHX, cycloheximide; HPG, L- homopropargylglycine; Tyr-TUB, tyrosinated tubulin. Bar, 5 µm. (**C**, **D**) Primary motoneurons (C) and quantification (D) of newly synthesized proteins (HPG, green) in WT and NKO somas stained with Tyr-TUB (grey). Data represent the mean ± SD of 3-4 mice (49-100 somas per culture). Statistical significance was determined by Dunnett’s multiple comparison test. Bar, 20 µm. (**E**) Volcano plot of the interactome of CLUH in HeLa cells. Green dots denote proteins involved in translation and RNA binding; red dots denote interactors of CLUH decreased in NKO axons. (**F**) KEGG pathway analysis of the interactome of CLUH in HeLa cells. Analysis was done using the EnrichR webtool.

If the lack of essential translational components distally underlies the axonal translation and GC defects in NKO motoneurons, the expression of depleted proteins could be beneficial. ABCE1 is the most downregulated protein related to translation in axons (Fig. 6F, G) and was enriched in the CLUH immunoprecipitated (Fig. 7E), and therefore caught our attention. ABCE1 is an essential regulator of initiation and termination of translation, recycling and quality control of ribosomes (*37*). Remarkably, transfected FLAG-tagged ABCE1 localizes to GCs, with a pattern very similar to that of overexpressed CLUH (Fig. S9C). We therefore expressed ABCE1 in NKO neurons and monitored axonal translation and the size of NKO GCs. Strikingly, expression of ABCE1 restored axonal protein synthesis (Fig. 8A, B) and the size of GCs of NKO motoneurons (Fig. 8C, D). In conclusion, we identify ABCE1 as a crucial translational component in axons, which decreases in axons upon CLUH loss and acts downstream of CLUH.

**Fig. 8:**
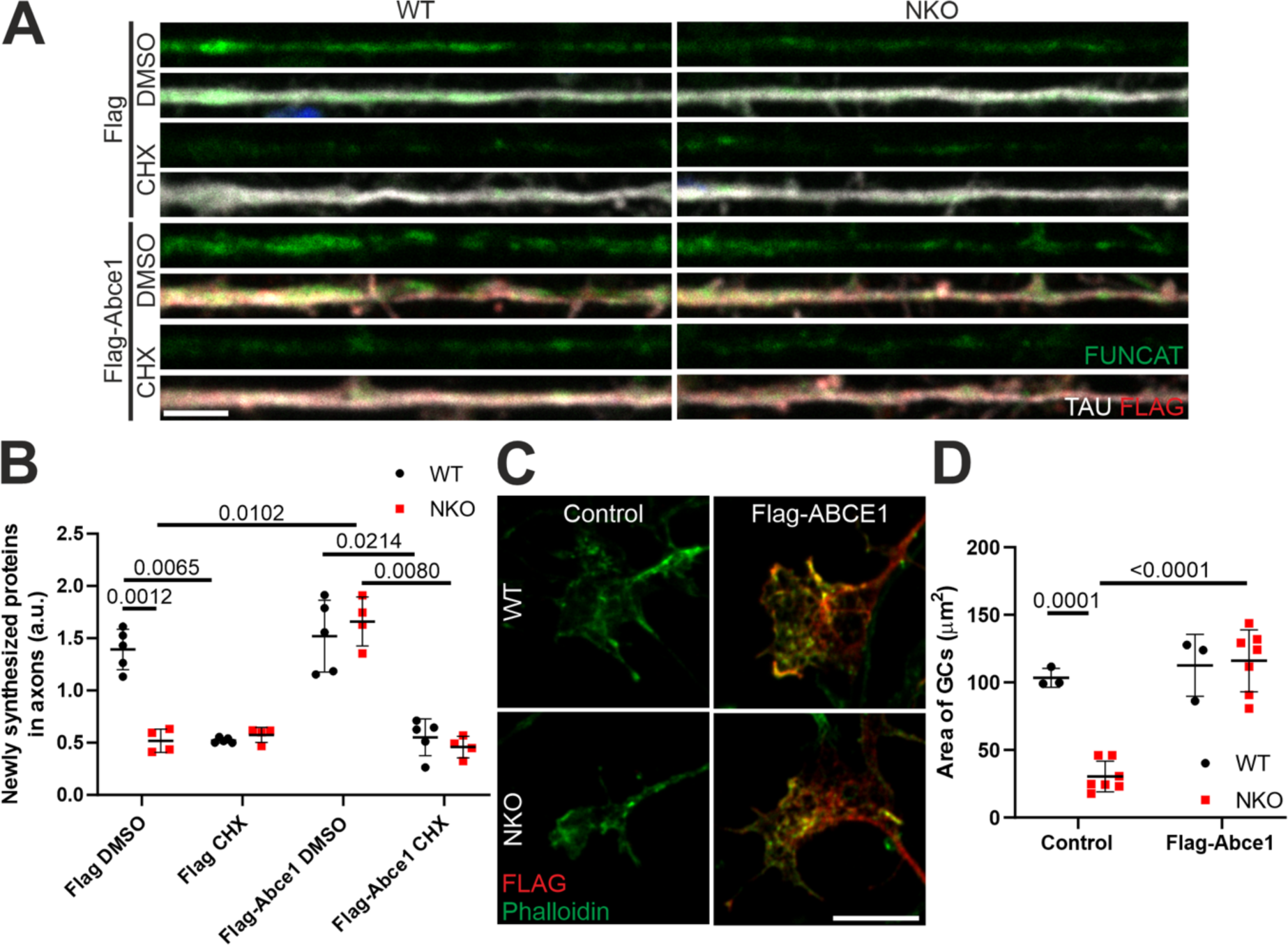
ABCE1 restores translation and GCs size in NKO axons (**A, B**) Single confocal planes (A) and quantification (B) of newly synthesized proteins (HPG, green) revealed by the FUNCAT assay in WT and NKO axons transfected with Flag or Flag- Abce1 (red) and stained with TAU (grey). Data represent the mean ± SD of 4-5 mice (10-22 axons per culture). Statistical significance was determined by Dunnett’s multiple comparison test. CHX, cycloheximide; HPG, L-homopropargylglycine; Tyr-TUB, tyrosinated tubulin. Bar, 5 µm. (**C, D**) Single confocal planes (C) and quantification (D) of the GCs area of GCs of control and FLAG- Abce1 (red) transfected motoneurons stained for Actin (Phalloidin, green). Data represent the mean ± SD of 3-7 cultures (17-34 GCs per culture). Statistical significance was determined by Dunnett’s multiple comparison test. Bar, 10µm.

## Discussion

Highly polarized neurons such as spinal motoneurons depend on mRNA transport and localized translation for development and survival, and often degenerate in neurological conditions caused by mutations in RBPs implicated in these processes (*38*). Here, we demonstrate a functional role of CLUH in peripheral axonal compartments. CLUH ablation in murine neural precursors leads to progressive motor dysfunction and degeneration of axons in the sciatic nerves while sparing the cell bodies. By using spinal motoneurons as an *in vitro* model system, we show that CLUH is essential to preserve the axonal pool of its target mRNAs and the respective proteins in axonal mitochondria, to maintain the ATP levels in distal mitochondria, and to sustain axonal growth. Surprisingly, CLUH has a critical function in supporting the translational capacity of axons.

By comparing the effect of CLUH depletion in neuronal somas versus axons or GCs, we found that lack of CLUH affects the abundance of targets mRNAs and their encoded proteins in neurites more than in the neuronal cell bodies. mRNA trafficking assays, using *Atp5a1* and *Mdh2* as exemplary targets, argued against a mere role of CLUH in transporting these transcripts along axons as the cause for their reduced axonal levels. The tagged mRNAs travelled along WT and NKO axons for the same time and with similar features in respect to directed versus oscillatory movement or confined localization. In further agreement with our conclusion is the observation of a decreased axonal abundance of the *Pink1* mRNA, which is known to be transported via binding to SYNJ2 and SYNJ2BP (*5*). The stability of mRNAs is *per se* an important determinant of neurite localization. Indeed, CLUH target mRNAs are depleted of destabilizing factors that render less likely that an mRNA reaches the peripheral regions of a neuron (*39*). 207 out of 229 putative CLUH targets contain less than 9 A-U rich elements in the 3’ UTR and only 26 out 229 are m^6^A (*N*^6^-methyladenosine)-enriched (*39*). Since we previously showed that the targets are subjected to faster decay in absence of CLUH in cellular models (*16, 19*), one supposition is that CLUH acts as a stabilizing factor during the mRNA life cycle. However, several lines of evidence strongly support the hypothesis of mRNA destabilization resulting from a translational defect. We found a distinct axonal signature in NKO axons, characterized by depletion of ribosomal components and translational regulators, in agreement with impaired axonal translation. We show that the CLUH interactome is enriched of ribosomal proteins and other translation components. Furthermore, other studies found CLUH in proximity to mitochondrial proteins before import, to mitochondrial targeting peptides, and to mitochondrially associated ribosomes (*40–42*).

We show that re-expression of ABCE1 in NKO axons was sufficient to rescue the translational defect and the GC size of NKO axons. ABCE1 is a fundamental player coupling translation termination and splitting of the two ribosomal subunits to translation initiation (*37*). By allowing the translational cycle to resume, ABCE1 counteracts the removal of ribosomes and associated mRNAs by ribophagy (*43*). Regulation of axonal translation occurs mainly at the level of initiation, and failure to activate translation at the right time and location, or to recycle ribosomes after a translational cycle could be especially detrimental. Our results are reminiscent of the beneficial role of ectopic ABCE1 expression in *Drosophila* to reverse mitochondrial aggregation caused by *Pink1* downregulation in the muscle. In this context, stalled ribosomes on damaged mitochondria lead to polyubiquitination of ABCE1 by NOT4. ABCE1 acts as a signal to initiate mitophagy by recruiting autophagic receptors to the mitochondria (*44*). In agreement with this role, ABCE1 overexpression also inhibits the accumulation of longer forms of respiratory chain proteins generated by co-translational extension upon mitochondrial dysfunction (*45*).

Strikingly, the *Drosophila* CLUH orthologue *clueless* interacts genetically with *Pink1* (*46*), and suppresses phenotypes of *Pink1* mutants when overexpressed (*46, 47*). Thus, PINK1 and CLUH may be involved in parallel pathways that initiate mitophagy upon different types of translational defects on mitochondrial surface. We previously found that mitophagy was affected in cells lacking CLUH (*48*). In addition, in primary hepatocytes CLUH was found together with target mRNAs in granules that were delivered to the lysosomes upon starvation (*48*). Thus, more experiments are needed to understand whether the lack of CLUH triggers some form of ribophagy, depleting axons of mRNAs for mitochondrial proteins and of ribosomal components, and whether mitophagy is hampered in CLUH-deficient axons.

Recent studies have shown that mitochondrial proteins are highly translated in axons (*10*), providing a partial explanation for the very pronounced translational defects of NKO axons. In dendrites, mitochondria act as reservoirs to provide the energy required for localized translation (*49*), thus the mitochondrial dysfunction caused by loss of CLUH may exacerbate defects in axonal translation. The pathogenic relevance of impaired mitochondrial respiration is supported by the ability of uridine supplementation to rescue the GC defects of NKO motoneurons. Uridine auxotrophy upon respiratory deficiency is traditionally linked to the mitochondrial enzyme dihydroorotate dehydrogenase (DHODH) that converts dihydroorotate in orotate, a step of pyrimidine biosynthesis. DHODH uses coenzyme Q as electron acceptor and is therefore dependent on a functional respiratory chain (*32*). Notably, DHODH is a putative CLUH target mRNA (*11*). A recent study has identified a novel role of uridine, as a precursor of ribose-1- phosphate, which can be converted to glycolytic intermediates (*35*). By fueling ATP production in the cytosol, uridine supplementation may therefore compensate a mitochondrial dysfunction by several mechanisms, reconciling previous studies showing a beneficial role of uridine to support neuronal survival under conditions of glucose deprivation (*50*).

Taken together, we identify CLUH as crucial player for the expression of mitochondrial proteins in axons and show the physiological role of CLUH to prevent peripheral neuropathy. The central and spinal motoneurons with long axons that show pathology upon CLUH depletion are affected also upon mutations in other RPBs, such as SMN, FUS, and TDP43 (*51*), which cause motoneuron disease and lead to defects in axonal translation and mitochondrial dysfunction. It will be interesting to explore if dysfunctional CLUH plays any role in the pathogenesis of these neurodegenerative diseases.

## Materials and Methods

### Mouse lines

All animal procedures were carried out in accordance to European (EU directive 86/609/EEC), national (Tierschutzgesetz) and institutional guidelines and were approved by local authorities (Landesamt für Natur, Umwelt und Vebraucherschutz Nordrhein-Westfalen, Germany). Animals were maintained in the CECAD Research Center, University of Cologne, Germany, in individually ventilated cages at 22°C (± 2°C) and a relative humidity of 55% (± 5%) under 12-h light cycle on sterilized bedding (Aspen wood, Abedd, Germany) and with access to sterilized commercial pelleted diet (Ssniff Spezialdiäten GmbH) and acidified water *ad libitum*. Previously described Cluh^fl/fl^ mice (*16*) were crossed with a nestin-Cre transgenic mice line (*23*) to generate a neural Cluh^fl/fl^ Cre^wt/tg^ (NKO) mice. Mice were kept on a pure C57/BL6N. Cluh^fl/fl^ Cre^wt/wt^ (WT) and Cluh^wt/wt^ Cre^wt/tg^ (Cre) mice were used as controls. Mice used for behavioral tests were analyzed by sex, tissues for histology and embryos were used independent of the sex. Animals were killed by CO2 inhalation to collect organs and pregnant females were killed by cervical dislocation to collect embryos. When required, mice were anesthetized with 20 mg xylazine/100 mg ketamine per kg of body-weight, and intracardially perfused with 4% paraformaldehyde (Sigma).

### Walking beam and rotarod tests

Animals were trained to cross a 90 cm long wide beam for 2 days. The training included the acclimatization from a 3 cm to a 1 cm wide beam and 10 min of rest between the two sessions. On the third day, animals were tested 3 times to cross the 1 cm wide beam. The number of slips and time to cross the beam were quantified. After 2 days of rest, mice were tested using a Rotarod apparatus (TSE system). Mice were trained for 5 min to acclimatize on the accelerating rod. The latency to fall from the apparatus was tested 3 times with a resting period of 15 min.

### Nerve conduction

CMAP amplitudes were recorded using a PowerLab single acquisition setup (ADInstruments, Grand Junction). After anesthesia, the sciatic nerves were stimulated using needle electrodes and the amplitude of the CMAP was recorded using recording needle electrodes into the hind paw as previously described (*52*).

### RNA extraction and transcriptomics

Freshly dissociated sciatic nerves from 5 months old mice were snapped frozen in liquid nitrogen and stored at -80°C. The RNA from sciatic nerves was isolated using the RNeasy Microkit (Qiagen) according to the instructions of the manufacturer. Samples were processed by the Cologne Center for Genomics. Due to low amount of input material, pre-amplification using the Ovation RNASeq System V2 was performed. Total RNA was used for first strand cDNA synthesis, using both poly(T) and random primers, followed by second strand synthesis and isothermal strand-displacement amplification. For library preparation, the Illumina Nextera XT DNA sample preparation protocol was used, with 1 ng cDNA input. After validation (Agilent 2200 TapeStation) and quantification (Invitrogen Qubit System) all six transcriptome libraries were pooled. The pool was quantified by using the Peqlab KAPA Library Quantification Kit and the Applied Biosystems 7900HT Sequence Detection System. The pool was sequenced on an Illumina NovaSeq6000 sequencing instrument with a PE100 protocol. RNA-seq was performed with a directional protocol. Quality control, trimming, and alignment were performed using the nf-core 56 RNA-seq pipeline (v3.0) (https://doi.org/10.5281/ZENODO.1400710). The reference genome sequence and transcript annotation used were Mus musculus genome GRCm39 from Ensembl version 103.Differential expression was analyzed in R version 4.1.2 (https://www.R-project.org/) with DESeq2 v1.34.0 (*53*) to make pairwise comparisons between groups. Log Fold Change shrinkage estimation was performed with ashr (*54*). Genes were considered as candidates to be differentially expressed if they had a minimum coverage of 10 reads in 6 or more samples from each pairwise comparison. Genes were significant if they met the following criteria: p-value < 0.05, q-value < 0.05 and fold change < -0.50 or > 0.50. Gene ontology analysis of significantly enriched genes was performed using the EnrichR webtool (*55*).

### Histology

For free-floating staining, 30 μm sections of fixed spinal cords were cut using a vibratome (VS1000, Leica), permeabilized and blocked with 0.4% Triton X-100 (Sigma) and 10% fetal bovine serum (FBS, Sigma) for 1 h and incubated with polyclonal goat anti-ChAT (AB144P, Millipore) in 5% FBS O/N. Then samples were incubated with donkey anti-goat IgG Alexa Fluor 488 (A11055, Thermo Fisher) for 2 h. Nuclei were counterstained using DAPI and mounted using ProLong Gold Antifade Reagent (Cell Signaling). For NMJ staining, freshly dissected transverse abdominal (TA) muscles were fixed with 4% PFA for 20 min, incubated in 0.1 M glycine for 30 min on an orbital shaker, permeabilized with 1% Triton X-100, blocked with 5% Bovine Serum Albumin (BSA, Sigma) for 2 h, incubated with primary antibody rabbit monoclonal anti-NEUROFILAMENT N (C28E10, Cell Signaling) in 2.5% BSA at 4°C ON, then with the following antibodies in 5% BSA for 1 h: donkey anti-rabbit IgG Alexa Fluor 488 (A21206, Thermo Fisher) and α-Bungarotoxin Alexa Fluor 594 (B13423, Thermo Fisher). Samples were mounted using Fluorsave reagent (Calbiochem).

Hindlimbs of E13.5 embryos were stained using a modified protocol from (*56*). Embryos were fixed in 4% PFA ON, permeabilized with 0.2 % TritonX-100/20 % DMSO at 37°C ON and then in 0.1% Tween-20/0.1% TritonX-100/0.1% Deoxycholate/0.1% NP40/20% DMSO at 37°C ON. Embryos were blocked in 0.2% TritonX-100/10% DMSO/6% goat serum at 37°C for 2 days and then incubated with mouse monoclonal anti-NF-M primary antibody (2H3, DSHB) in 0.2% Tween-20 with 10ug/ml heparin/5% DMSO/3% goat serum at 37°C for 3 days, followed by the incubation with secondary antibody goat anti-mouse IgG Alexa Fluor 488 at 37°C for 2 days. Images were acquired using the HC PL APO CS2 20x/0.75 DRY and HC PL APO CS 40x/0.85 DRY objectives of an SP8 confocal microscope (Leica).

### Semithin sections and electron microscopy

Sciatic nerves from perfused animals were post-fixed in 2% glutaraldehyde (Sigma) in 0.1 M cacodylate buffer (Sigma). Sciatic nerves were cut distally and stained with 1% toluidine blue. Semithin sections were imaged using the Plan-Apochromat 20X/0.8 objective of an Axio-Imager M2 microscope equipped with Apotome 2 (Zeiss). The number of axons was automatically segmented and quantified using the Trainable Weeka Segmentation plugin of ImageJ (National Institutes of Health, Bethesda). Degenerating axons were manually quantified as dark spots and normalized by the surface of the cross section. For ultrastructural studies, sciatic nerves were embedded in Epon and imaged as previously described (*16*), using a transmission electron microscope (JEOL JEM2100PLUS) provided with a GATAN OneView camera. The morphology of mitochondria was manually assessed by an experimenter blind to the experiment.

### DNA constructs

Mito-GFP (*57*), pcDNA-AT1.03 and pcDNA-mitoAT1.03 (*29*), pcDNA3.1-FLAG-Abce1 (*58*) (kindly provided by R.S. Hedge) were previously described. Full-length human CLUH and mouse *Cluh* coding sequences cDNA were cloned in pcDNA3 and pmCherry-N1 (Clontech), respectively, using HindIII and EcoRI restriction sites. For mRNA trafficking, target mRNAs were tagged with 24 repeats of the MS2V5 sequence behind their 3’UTR. To achieve these constructs the HaloTag-bActinCDS-bActinUTR-MS2V5 plasmid (Addgene #102718) (*59*) was used as a basis for the cloning. An initial restriction digest using NotI and AgeI removed all tags and the *ActB* mRNA, leaving only the MS2V5 sequence. *ActB*, *Mdh2* and *Atp5a1* mRNA were cloned back into the backbone by Gibson Assembly, using the Addgene Plasmid #102718 as a template for *ActB* and mouse cDNA as a template for *Mdh2 (*ENSMUST00000019323) and *Atp5a1* (ENSMUST0000002649). For the reporter plasmid NLS-HA-stdMCP-stdHalo (Addgene # 104999) (*60*) was used.

### Cell lines and cell culture

Immortalized MEFs were previously described (*11*). MEFs were electroporated using p3 primary cell 4d-nucleofector x kit L (V4XP-3024, Amaxa) using Nucleofector I (Amaxa). Spinal cords were dissected from E13.5 embryos and primary motoneurons were isolated as previously described (*61*). Motoneurons were plated on coverslips coated with 20 μg/ml poly-D-lysine (Sigma) and 0.1 μg/ml laminin (Sigma) at a density of 15,000 cells/cm^2^ and grown in Neurobasal medium (Thermo Fisher) supplemented with 2% B-27 (Thermo Fisher), 1% glutamine (Thermo Fisher), 1% Pen/Strep (Thermo Fisher), 250 μg/ml Amphotericin B (Promocell), 1 μM cytosine arabinoside (Sigma), 10 ng/ml brain-derived neurotrophic factor (BDNF, Peprotech), 10 ng/ml ciliary neurotrophic factor (CNTF, Peprotech), and 10 ng/ml glia cell line-derived neurotrophic factor (GDNF, Peprotech). The medium was supplemented with 50 μg/ml uridine where indicated. Cells were cultured in a humified incubator with 5% CO2 at 37°C. For neurons grown in Boyden chambers (1.0 μm pore size, Falcon), the medium contained in the bottom compartment was supplemented with 5 times the concentration of BDNF, CNTF and GDNF. Neurons were transfected with Lipofectamine 2000 (Thermo Fisher) according to (*62*).

### Immunocytofluorescence

Cells were fixed with 4% formaldehyde (Sigma) for 20 min, permeabilized with 0.1% Triton X- 100 (Sigma) for 10 min, blocked with 10% goat serum for 1 h and incubated with primary antibodies in 1% goat serum O/N. The following antibodies were used: monoclonal rabbit anti- ABCE1 (ab32270, Abcam), polyclonal rabbit anti-CLUH (NB100-93306, Novus Biologicals), monoclonal mouse anti-Cherry (677702, Biolegend), monoclonal rat anti-CLASP2 (MAB9738, Abnova), mouse monoclonal anti-FLAG (F3165, Sigma), polyclonal rabbit anti-IQGAP1 (2217- 1-AP, Proteintech), polyclonal rabbit anti-MAP2 (4542, Cell Signaling), polyclonal rabbit anti-RFP (600-401-379S, Rockland), polyclonal rabbit anti-RPL14 (14991-1-AP, Proteintech), polyclonal rabbit anti-RPS8 (ab201454, Abcam), polyclonal rabbit anti-SYNAPSIN1/2 (106002, Synaptic Systems), monoclonal mouse anti-TAU (sc-390476, Santa Cruz Biotechnology), polyclonal rabbit anti-TOM20 (sc-114115, Santa Cruz Biotechnology), monoclonal mouse anti-β-TubulinIII (T8660, Sigma), monoclonal mouse anti-Acetylated-Tubulin (T6793, Sigma), monoclonal mouse anti-Tyrosinated-Tubulin (T9028, Sigma). Samples were further incubated with secondary antibodies in 1% goat serum for 2 h. Actin was stained using Alexa Fluor 555 Phalloidin or Alexa Fluor 488 Phalloidin (Thermo Fisher). The following antibodies were used: goat anti-mouse IgG Alexa Fluor 488 (11029, Thermo Fisher), goat anti-mouse IgG Alexa Fluor 594 (A11005, Thermo Fisher), goat anti-mouse IgG Alexa Fluor 647 (A21236, Thermo Fisher), goat anti-rabbit IgG Alexa Fluor 488 (A11034, Thermo Fisher), goat anti-rabbit IgG Alexa Fluor 546 (A11035, Thermo Fisher), goat anti-rat IgG Alexa Fluor 546 (A11081, Thermo Fisher).

Nuclei were counterstained using DAPI and the coverslips were mounted using Fluorsave reagent (Calbiochem). Images were acquired using the HC PL APO CS2 20x/0.75 DRY and HC PL APO CS 40x/0.85 DRY objectives of an SP8 confocal microscope (Leica). The morphology of mitochondria was quantified using the macro Mitomorph (*63*) for ImageJ. The intensity of fluorescent signal was quantified using the mean gray-value function of ImageJ. The length of axons in Boyden chambers was quantified using the AxonTracer plugin for ImageJ (*64*).

### RNA in situ hybridization

RNA in situ hybridization following the instruction of the RNAscope 2.5 HD Fluorescent Reagent Kit (Advanced Cell Diagnostics). Neurons were fixed in 10% formalin, dehydrated and rehydrated in a gradient of 50-70-100% ethanol of 5 min, incubated with 1% Tween-20 for 15 min and with Protease III for 10 min. Samples were then incubated with probes designed by the manufacturer to detect the corresponding mRNAs: *Atp5a1* (459311, Advanced Cell Diagnostics), *Pink1* (524081, Advanced Cell Diagnostics) *Actb* (316741, Advanced Cell Diagnostics). Signal was amplified according to the instruction of the manufacturer and samples were subjected to the immunofluorescence protocol described previously. Axons were straightened using the Straighten plugin of ImageJ and the number of mRNA dots was automatically quantified using a macro developed by the CECAD imaging facility.

### FUNCAT assay

Newly synthetized proteins were labelled using the Click-iT HPG Alexa Fluor488 Protein Synthesis Assay Kit (Thermo Fisher). Neurons were incubated with 0.1 mM L-homopropargylglycine (Thermo Fisher) for 30 min in L-methionine-free medium. Treatment with 0.1 mg/ml cycloheximide for 3 h was used as a negative control. Neurons were fixed in 4% FA for 15 min, permeabilized with 0.1% Triton X-100 for 10 min, incubated with Alexa Fluor azide 488 for 45 min and then subjected to the immunofluorescence protocol previously described. Axons were selected and the intensity of the fluorescence was measured using the multimeasure function of ImageJ.

### Apoptosis assay

Neurons were fixed in 4% FA for 15 min, permeabilized with 0.1% Triton X-100 for 10 min and stained using the In-Situ Cell Death Detection Kit, Fluorescein (Roche). Nuclei were counterstained using DAPI. Apoptosis was evaluated as the number of TUNEL^+^ cells respect to total DAPI^+^ nuclei.

### Live imaging

Live imaging experiments were performed using the LSM980 Airyscan 2 confocal microscope (Carl Zeiss Microscopy) equipped with the C-Apochromat 40X/1.20 W Korr or the Plan-Apochromat 63x/1,4 Oil DIC objective.

#### Measurement of ATP

Neurons were grown in complete Neurobasal medium or in complete Neurobasal medium supplemented with 9.0 g/L of galactose, as indicated. Neurons transfected with pcDNA-AT1.03 or pcDNA-mitoAT1.03 were imaged using the C-Apochromat 40X/1.20 W Korr objective of an Airy Scan confocal (Zeiss). Samples were treated with 0.01 mg/ml oligomycin (Sigma). FRET images were quantified using the fluorescent intensity function of ImageJ.

#### Mitochondrial potential

Neurons were grown in Boyden chambers as previously described. Membranes were incubated with 25 nM Tetramethylrhodamine, methyl ester (TMRM, Sigma) for 30 min. Samples were treated with 0.01 mg/ml oligomycin and with 2 μM carbonyl cyanide m-chlorophenyl hydrazone (CCCP, Sigma). Images from the axonal compartment were acquired 1 per min. The fluorescent signal from mitochondria was measured using the multimeasure function of ImageJ and normalized to the initial value.

#### Trafficking of mRNA

Primary motoneurons were co-transfected at DIV 4 with a 25:1 ratio of mRNA-MS2 and MCP-Halo. 24 h later the cells were incubated with a final concentration of 16 nM Halo-Ligand JF 549 (Promega, GA1110) for 30 min, followed by one replacement of the medium. Neurites with visible mRNA dots were selected and a timelapse was taken with three z-slices per frame, one frame every 0.6 seconds and for 90 seconds overall. Movies were analyzed on the one hand by Kymographs which were generated from the maximum projected time lapses using ImageJ. In the Kymographs mRNA tracks were manually traced and the tracks were analysed using the step-3 measure of the Kymolyzer macro (*65*) On the other hand movies were analyzed with the automatic tracking module of the NIS-element AR software (Nikon). Only tracks with more than 20 frames were considered. A custom-made add-on Matlab script was developed to analyze the type of movement and directionality of mRNA trajectories in neurites. In particular, to define the type of movement of each particle the mean square displacement (MSD) was calculated as follows: MSD(τ) = < (x(t+τ)–x(t))2+(y(t+τ)–y(t)) 2 > (Eq1) where x and y are the coordinates of the mRNA along the neurite, t and τ are the absolute and lag times, respectively, and the brackets represent the time average. This calculation was performed for τ = 25% of the total time of the trajectory (*66*). The MSD data were fitted with an anomalous diffusion model: MSD = Aτα+B (Eq2) where A depends on the motion properties of the particle, B is the residual MSD, and the coefficient α is an indication of the particle motion-type (*67*).

Trajectories were classified as actively driven (α > 1.5), diffusive (0.9 < α <1.1) or confined (α < 0.5). Finally, to define the motion type based on the lateral displacement of mRNAs the net displacement (ND) and lateral maximal displacement (LMD) were measured. ND is defined as the difference in x coordinates of the first point and the last point of the trace. Lateral maximal displacement (LMD) is defined as the difference between the first point and the most distal point during the trace. mRNAs with ND > 5 µm are defined directed; mRNAs with ND < 5 µm and LMD < 1 µm were defined stationary while particles with ND < 5 µm and LMD > 1 µm were defined oscillatory (*68*).

### Real-time PCR

RNA from primary neurons was isolated using TRIzol (Thermo Fisher). cDNA was retrotranscribed from 1 μg of RNA using the SuperScript First-Strand Synthesis System (Life Technologies). Real-time PCR was performed using SYBR Green Master Mix (Applied Biosystems) with the Quant Studio 12K Flex Real-Time PCR System thermocycler (Applied Biosystems). The following primers were used: ChAT forward 5’- CCTGCCAGTCAACTCTAGCC-3’, reverse 5’-TACAGAGAGGCTGCCCTGAG-3’; Hb9 forward 5’-CCAAGCGTTTTGAGGTGGC-3’, reverse 5’- GGAACCAAATCTTCACCTGAGTCT-3’; Gapdh forward 5′-AGGTCGGTGTGAACGGATTTG-3′, reverse 5′-TGTAGACCATGTAGTTGAGGTCA-3′. The fold enrichment was calculated using the 2^(−ΔΔCt)^ formula. To quantify the mitochondrial DNA, the following primers were used: *mt-Co1* forward 5′-TGCTAGCCGCAGGCATTACT-3′, reverse, 5′- CGGGATCAAAGAAAGTTGTGTTT-3′; *Rmrp* forward 5′- GCCTACACTGGAGTCGTGCTACT-3′, reverse 5′-CTGACCACACGAGCTGGTAGAA-3′.

### Protein isolation and western blot

2 million motoneurons were plated and grown for the indicated days. Proteins were extracted using RIPA buffer (1% sodium cacodylate, 50 mM Tris-HCl, 150 mM NaCl, 5 mM EDTA, 1% Triton-X-100) and protease inhibitor cocktail (Sigma). For tissues, RIPA buffer was supplemented with 0.1% SDS. Proteins were quantified using standard Bradford (Bio-Rad) assay and subjected to SDS-PAGE and transferred on PVDF membranes. The following antibodies were used: monoclonal mouse anti-pan-actin (MAB1501R, Millipore) and polyclonal rabbit anti-CLUH (ARP70642_P050, Aviva).

### Immunoprecipitation of CLUH in HeLa cells followed by MS

HeLa were lysed in IP buffer (50 mM Tris-HCl, pH 7.4, 50 mM KCl, 0.1% Triton X-100 and freshly added protease inhibitor cocktail). Bradford assay was used to quantify the protein concentration and 400 μg of cell lysate was diluted in 250 μl of IP buffer and incubated with rabbit polyclonal anti-CLUH antibody (NB100-93306, Novus Biologicals) or rabbit IgG isotope control antibody (NB810-56910, Novus Biologicals) for 3 h in the cold room with constant agitation. Afterwards, the samples were incubated with 20 μl of prewashed Protein G Dynabeads (10003D, Thermo Fisher) with constant agitation in the cold room for 1 h. This was followed by 5 washes with IP buffer on a magnetic stand, the proteins were then heat eluted at 95°C for 5 min in 30 μl of SP3 lysis buffer (5% SDS in PBS). The supernatants were treated with 5 mM DTT (dithiothreitol) at 55°C for 30 min and then with 40 mM CAA (chloroacetamide) at RT for another 30 min in the dark. After centrifugation, supernatants were processed by the CECAD Proteomics Facility. All samples were analyzed on a Q-Exactive Plus (Thermo Scientific) mass spectrometer that was coupled to an EASY nLC 1000 or 1200 UPLC (Thermo Scientific). Samples were loaded onto an in-house packed analytical column (50 cm × 75 µm I.D., filled with 2.7 µm Poroshell EC120 C18, Agilent) that has been equilibrated in solvent A (0.1 % formic acid in water) and peptides were separated at a constant flow rate of 250 nL/min using a 50 min gradient followed by a 10 min wash with 95 % Solvent B (0.1% formic acid in 80% acetonitrile). The mass spectrometer was operated in data-dependent acquisition mode. MS1 survey scans were acquired from 300 to 1750 m/z at a resolution of 70,000. The top 10 most abundant peptides were isolated within a 1.8 Th window and subjected to HCD fragmentation at a normalized collision energy of 27%. The AGC target was set to 5e5 charges, allowing a maximum injection time of 110 ms.

Product ions were detected in the Orbitrap at a resolution of 35,000. Precursors were dynamically excluded for 10 s. All mass spectrometric raw data were processed with MaxQuant (version 1.5.3.8) using default parameters. Briefly, MS2 spectra were searched against the Swiss-Prot human fasta database containing canonical sequences (downloaded 08.26.2020), including a list of common contaminants. False discovery rates on protein and PSM level were estimated by the target-decoy approach to 1% (Protein FDR) and 1% (PSM FDR) respectively. The minimal peptide length was set to 7 amino acids and carbamidomethylation at cysteine residues was considered as a fixed modification. Oxidation (M) and Acetyl (Protein N-term) were included as variable modifications. The match-between runs option was enabled within replicate groups. LFQ quantification was enabled default settings. Student’s T-tests were calculated in Perseus (version 1.6.15.0) after removal of decoys and potential contaminants. Data were filtered for at least 4 out of 4 values in at least one condition. Remaining missing values were imputed with random values from the lower end of the intensity distribution using Perseus defaults. Proteins were considered significant interactors with a p-value < 0.05, q-value < 0.05 and fold change > 0.40. KEGG pathway analysis of significantly enriched proteins was performed using the EnrichR webtool (*55*).

### Proteomics of primary neurons cultured in Boyden chambers

Neurons were grown in Boyden chambers for 6 days. Axons were scraped and neurons were lysed with 4% SDS in 100 mM Hepes pH 8.5 and snap frozen into liquid nitrogen. Samples were thawed and proteins were reduced (10 mM TCEP) and alkylated (20 mM CAA) in the dark for 45 min at 45 °C. Then samples were heated to 10 min incubation at 70°C on a ThermoMixer (shaking: 550 rpm). For neuron samples, the protein concentration was determined using the 660 nm Protein Assay (Thermo Fisher Scientific, #22660) and 20 µg of protein was subjected to tryptic digestion. For axon samples, the total volume was utilized. Samples were subjected to an SP3-based digestion (Hughes et al., 2014). Washed SP3 beads (SP3 beads (Sera-Mag(TM) Magnetic Carboxylate Modified Particles (Hydrophobic, GE44152105050250), Sera-Mag(TM) Magnetic Carboxylate Modified Particles (Hydrophilic, GE24152105050250) from Sigma Aldrich) were mixed equally, and 3 µL (soma) and 1 µL (axon) of bead slurry were added to each sample. Acetonitrile was added to a final concentration of 50% and washed twice using 70 % ethanol (V = 200 µL) on an in-house made magnet. After an additional acetonitrile wash (V = 200µL), 5 µL (soma) or 0.5 µL (axon) digestion solution (10 mM HEPES pH = 8.5 containing 0.5µg Trypsin (Sigma) and 0.5µg LysC (Wako) per µL) was added to each sample and incubated ON at 37°C. Peptides were desalted on a magnet using 2 x 200 µL acetonitrile. Peptides were eluted in 10 µL 5% DMSO in LC-MS water (Sigma Aldrich) in an ultrasonic bath for 10 min.

Formic acid and acetonitrile were added to a final concentration of 2.5% and 2%, respectively. The protocol was automated using CyBio Felix (Analytik Jena) and Mosquito LV (SPT Labtech) liquid handler. Samples were stored at -20°C before subjection to LC-MS/MS analysis.

#### Liquid chromatography and mass spectrometry

LC-MS/MS instrumentation consisted of an Easy-LC 1200 (Thermo Fisher Scientific) coupled via a nano-electrospray ionization source to an Exploris 480 mass spectrometer (Thermo Fisher Scientific, Bremen, Germany). An in-house packed column (inner diameter: 75 µm, length: 20 cm) was used for peptide separation. A binary buffer system (A: 0.1% formic acid and B: 0.1 % formic acid in 80% acetonitrile) was applied as follows: Linear increase of buffer B from 4% to 27% within 40 min, followed by a linear increase to 45% within 5 min. The buffer B content was further ramped to 65% within 5 min and then to 95 % within 5 min. 95% buffer B was kept for a further 5 min to wash the column. Prior to each sample, the column was washed using 5 µL buffer A and the sample was loaded using 8 µL buffer A.

The RF Lens amplitude was set to 55%, the capillary temperature was 275°C and the polarity was set to positive. MS1 profile spectra were acquired using a resolution of 60,000 (at 200 m/z) at a mass range of 320-1150 m/z and an AGC target of 1 × 10^6^.

For MS/MS independent spectra acquisition, 48 equally spaced windows were acquired at an isolation m/z range of 15 Th, and the isolation windows overlapped by 1 Th. The fixed first mass was 200 m/z. The isolation center range covered a mass range of 357–1060 m/z. Fragmentation spectra were acquired at a resolution of 15,000 at 200 m/z using a maximal injection time of 22 ms and stepped normalized collision energies (NCE) of 26, 28, and 30. The default charge state was set to 3. The AGC target was set to 3e6 (900% - Exploris 480). MS2 spectra were acquired in centroid mode. FAIMS was enabled and used at a compensation voltage (CV) of -50 for all samples using an inner electrode temperature of 89°C and an outer electrode temperature of 99.5°C.

#### Data analysis

DIA-NN (Data-Independent Acquisition by Neural Networks) v 1.8 (*69*) was used to analyze data-independent raw files. Axon and soma samples were analyzed separately. The spectral library was created using the reviewed-only Uniport reference protein (Mus musculus, 17029 entries) with the ‘Deep learning-based spectra and RTs prediction’ turned on. Protease was set to Trypsin and a maximum of 1 miss cleavage was allowed. N-term M excision was set as a variable modification and carbamidomethylation at cysteine residues was set as a fixed modification. The peptide length was set to 7 – 30 amino acids and the precursor m/z range was defined from 340 – 1200 m/z. The option ‘Quantitative matrices’ was enabled. The FDR was set to 1 % and the mass accuracy (MS2 and MS1) as well as the scan window was set to 0 (automatic inference via DIA-NN). Match between runs (MBR) was enabled. The Neuronal network classifier worked in ‘double pass mode’ and protein interference was set to ‘Isoform IDs’. The quantification strategy was set to ‘robust LC (high accuracy)’ and cross-run normalization was defined as ‘RT-dependent’. The ‘pg’ (protein group) output (MaxLFQ intensities (*70*)) was further processed using Instant Clue (*71*) including a pairwise one-sample t-test utilizing the isolation and litter- batch derived fold change between NKO and WT. MitoCarta 3.0 and Uniprot-based Gene Ontology annotations were added and used for filtering. Volcano plots were generated using Prism 8.3.0 (GraphPad Software, LLC). Gene ontology analysis of significantly enriched proteins (p-value < 0.05; fold change as indicated in the main text) was performed using the EnrichR webtool (*55*).

### Software and statistical analysis

Templates for images or schemes were generated using Biorender. Prism 8.3.0 (GraphPad Software, LLC) was used for statistical analysis. In graphs, average and standard deviation (± SD) are reported, where each dot represents one independent experiment blind to the experimenters.

The number of experiments and samples per experiment are reported in figure legends. Normality was assessed using the Shapiro-Wilk test. p-values and statistical tests are reported in figures and figure legends.

## Acknowledgments

We thank the Cologne Center for Genomics and the CECAD proteomics, bioinformatics, imaging and in vivo research facilities for the technical assistance. We thank Peter Zentis for developing a macro to quantify RNAscope experiments. We thank Désirée Schatton, David Pla-Martin, and Paola Martinelli for the generation of constructs and preliminary data. We are grateful to Maximilian Thelen and Min Kye (University of Cologne) for support with neuronal cultures and Steffen Hermans (MPI for Biology of Ageing, Cologne) for the help with the rotarod test. We thank Désirée Schatton, Jan Riemer (University of Cologne) and Christian Frezza (University of Cologne) for suggestions and critical reading of the manuscript. Some reagents were kindly shared by Ramanujan Hedge (MRC, Cambridge), Sichen Shao (Harvard Med School, Boston) and Matteo Bergami (University of Cologne). This work was supported by the Deutsche Forschungsgemeinschaft (Project number 269925409).

## Contributions

M.Z. and E.I.R. conceived the project. M.Z., T.S., H.N., M.P., H.W., E.S. and T.U. performed and/or analysed experiments with the assistance from E.B. and L.W. E.I.R., M.Z., T.L. wrote the manuscript. E.I.R, T. L., H.C.L., B.W. and J-M. C. supervised the projects. All authors commented and approved the manuscript.

## Data and materials availability

All data are available in the main text or the supplementary materials.

## Supplementary Figures

**Supplementary Fig. 1:**
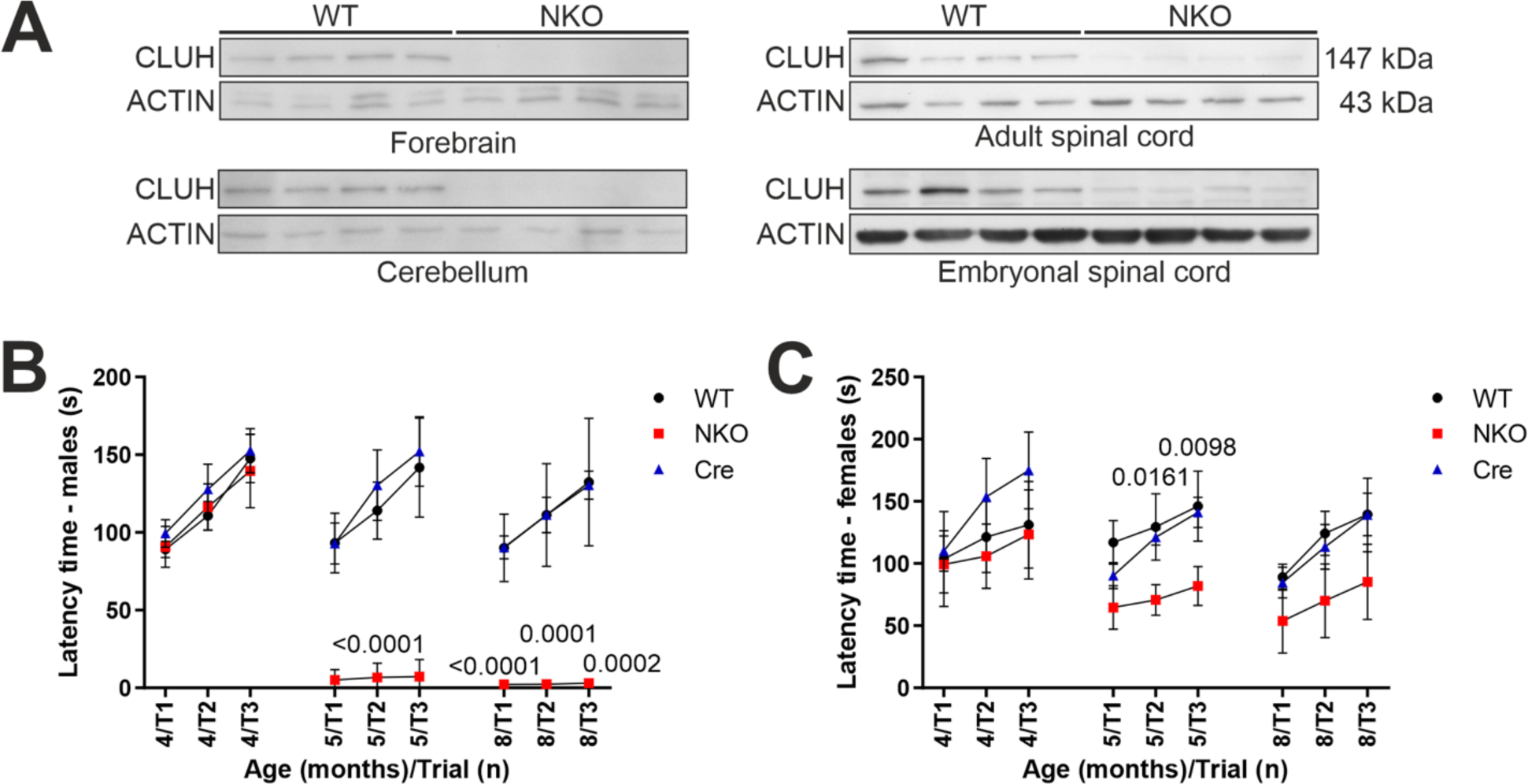
NKO mice display an impaired rotarod performance (**A**) Western blots depicting CLUH levels in forebrain, cerebellum and spinal cord of WT and NKO mice aged 5 months and spinal cord of WT and NKO E13.5 embryos. (**B, C**) Latency time to fall from a rotarod apparatus of males (B) and females (C) mice of the indicated genotypes. Data represent mean ±SD of 6-11 mice per group. Statistical significance was determined by Dunn’s and Dunnett’s multiple comparison tests.

**Supplementary Fig. 2:**
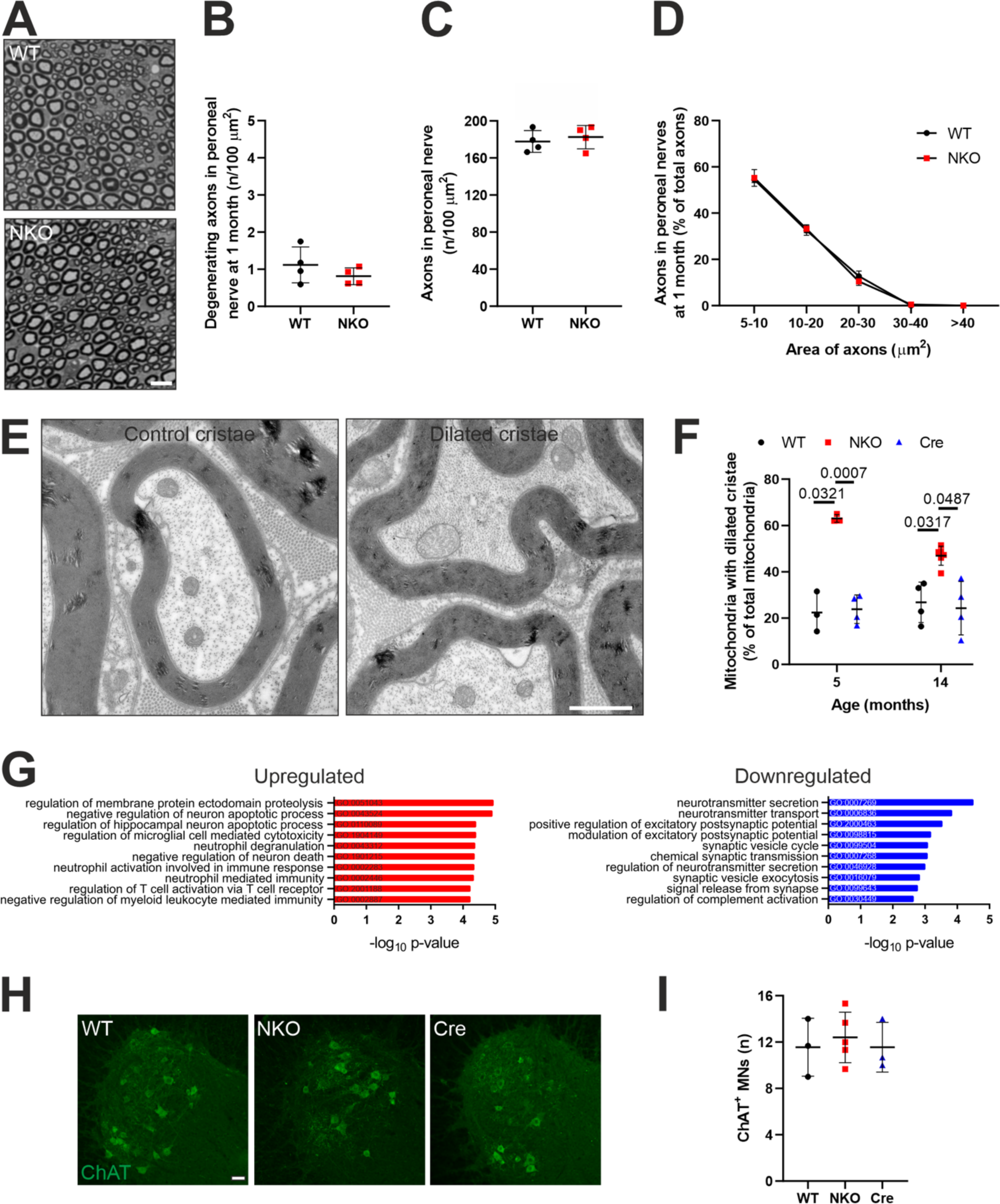
NKO motoneurons do not die *in vivo*, but show signs of axonopathy (**A**) Semithin sections of peroneal nerves of 1 month old mice. Bar, 10 µm. (**B**) Quantification of degenerating axons in peroneal nerves in experiments as in (A). Data represent mean ± SD of 4 mice. (**C**) Quantification of number of axons larger than 5 µm^2^ in peroneal nerves in experiments as in (A). Data represent mean ± SD of 4 mice. (**D**) Distribution of axons per area in peroneal nerves in experiments as in (A). Data represent mean ± SD of 4 mice. (**E, F**) Electron micrographs (E) and quantification (F) of the percentage of mitochondria with dilated cristae in axons of the tibial brach of the sciatic nerve of mice at 5 and 14 months of age. Data represent mean ± SD of 3-6 mice (55-219 mitochondria per mouse). Statistical significance was determined by Dunnett’s multiple comparison test. (**G**) Gene ontology biological process (GOBP) analysis of upregulated (left, red) and downregulated (right, blue) transcriptome changes in the sciatic nerves of 5 months old male NKO mice compared to WT mice. Analysis was done using the EnrichR webtool. (**H**) Anterior spinal cord stained for Choline Acetyltransferase (ChAT, green) in mice of indicated genotypes at 14 months of age. Bar, 50 µm. (**I**) Quantification of the number of motoneurons in experiments shown in (G). Data represent mean ± SD of 3-5 mice (4-6 sections per mouse).

**Supplementary Fig. 3:**
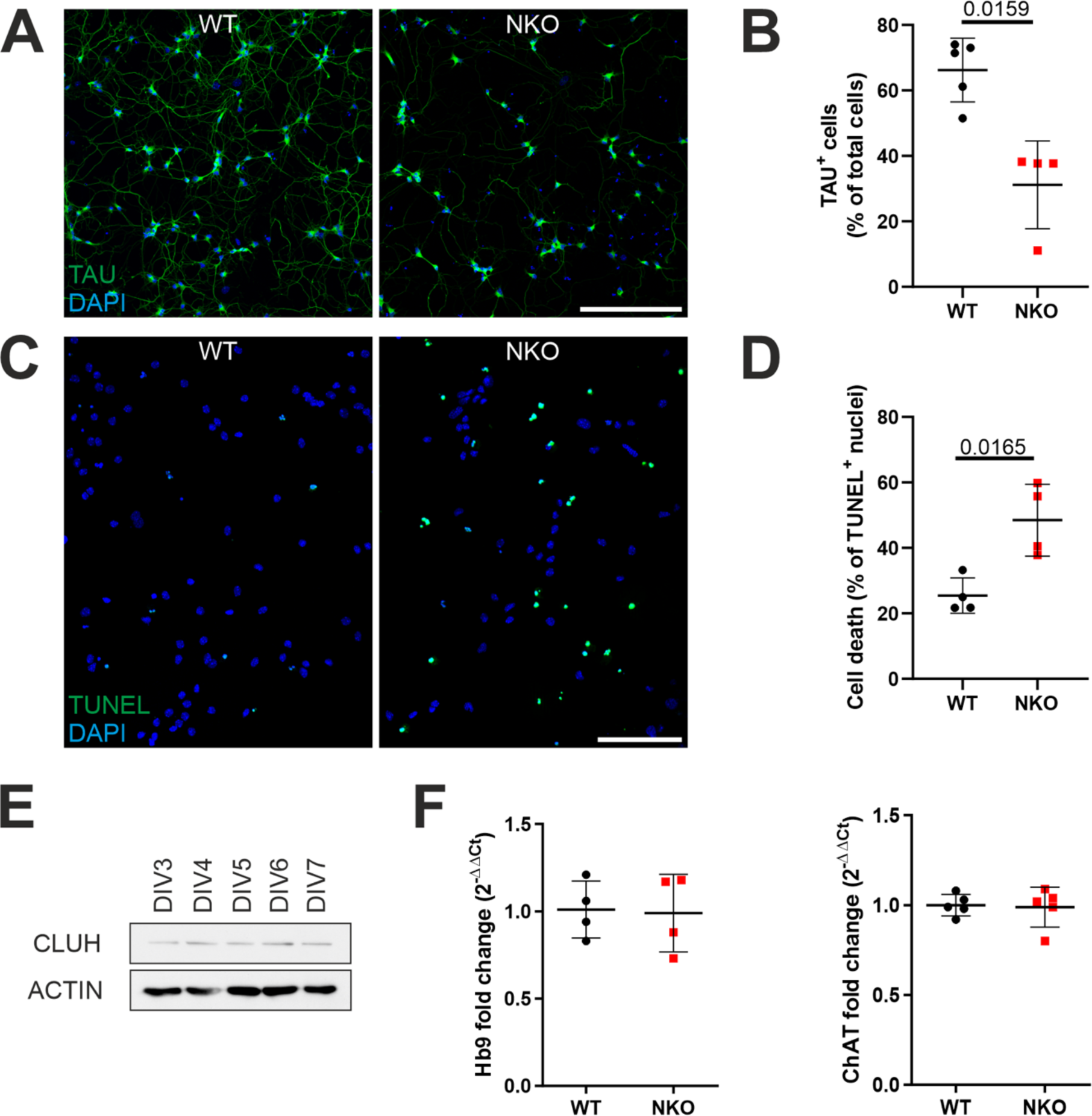
Phenotypes of NKO motoneurons (**A**) Primary spinal motoneurons at DIV 10 stained with TAU (green) and DAPI (blue). Bar, 50 µm. (**B**) Quantification of the number of TAU^+^ neurons in experiments as in (A). Data represent mean ± SD of 4-5 mice (20 fields per mouse). Statistical significance was determined by Mann-Whitney test. (**C**) Nuclei (DAPI, blue) and apoptotic nuclei stained using TUNEL assay (green) of primary motoneurons at DIV 10. Bar, 50 µm. (**D**) Quantification of apoptotic nuclei in experiments as in (C). Data represent mean ± SD of 4 mice (10 fields per mouse). Statistical significance was determined by Welch’s t test. (**E**) Western blot depicting CLUH levels in primary motoneurons from DIV 3 to DIV 7. (**F**) Quantification of the expression of Hb9 and ChAT in primary motoneurons at DIV 5. Data represent mean ±SD of 4-5 mice.

**Supplementary Fig. 4:**
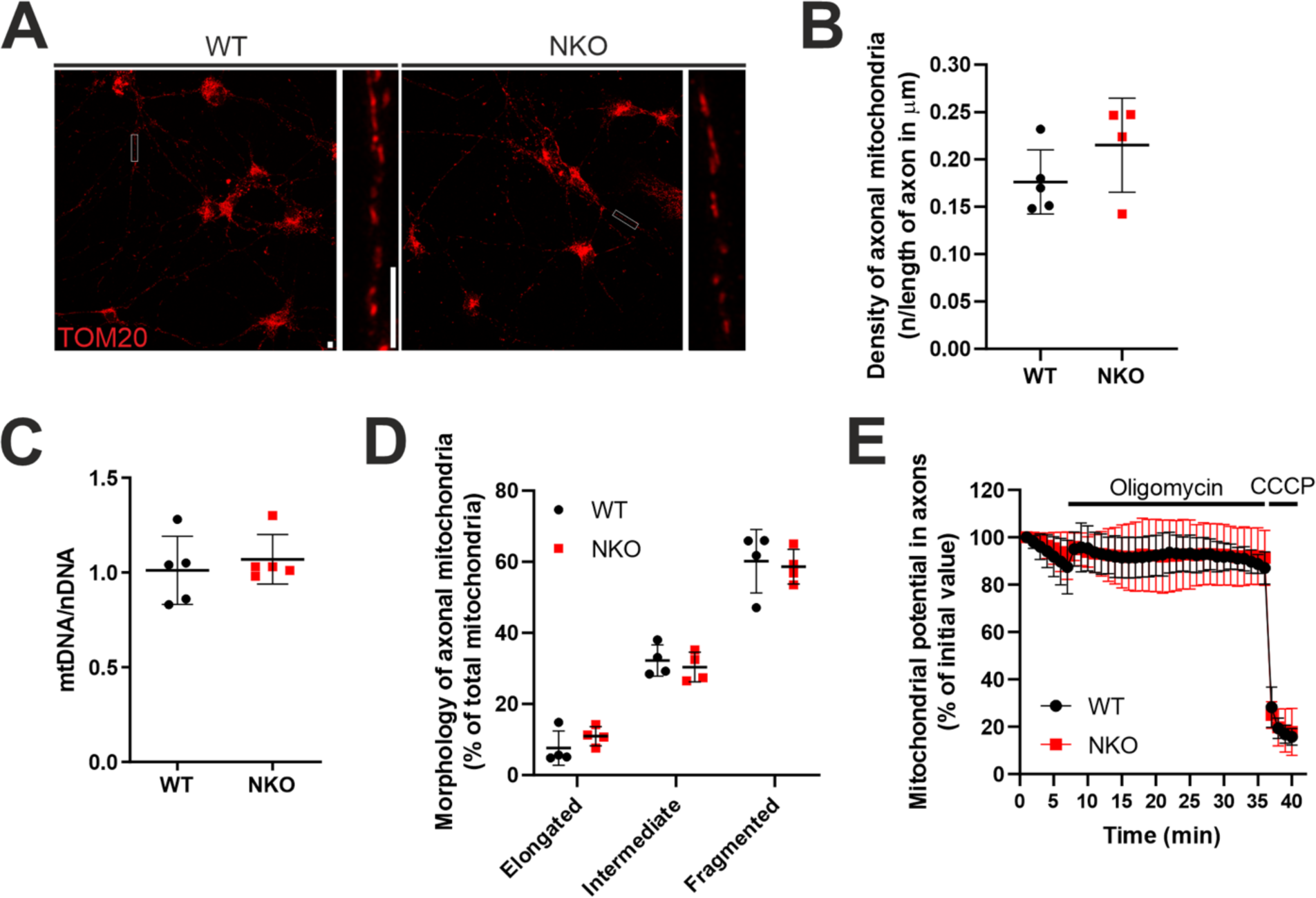
NKO mitochondria are comparable to WT mitochondria in respect to clustering, morphology and potential (**A**) Primary motoneurons stained with TOM20 (red). Representative axons are straightened and magnified on the right of each panel. Bars, 5 µm. (**B**) Quantification of the density of mitochondria in axons in experiments shown in (A). Data represent mean ± SD of 4-5 cultures (24-35 axons per culture). (**C**) Quantification of the abundance of mitochondrial DNA (mtDNA) to nuclear DNA (nDNA). Data represent mean ± SD of 5 motoneuronal cultures. (**D**) Quantification of the morphology of mitochondria in axons in experiments shown in (A). Data represent mean ±SD of 4 cultures (28-30 axons per culture). (**E**) Quantification of the mitochondrial potential of axons of primary motoneurons loaded with TMRM. Neurons were treated with oligomycin after 5 minutes and CCCP after 35 minutes. Data represent mean ± SD of 4 WT and 5 NKO cultures.

**Supplementary Fig. 5:**
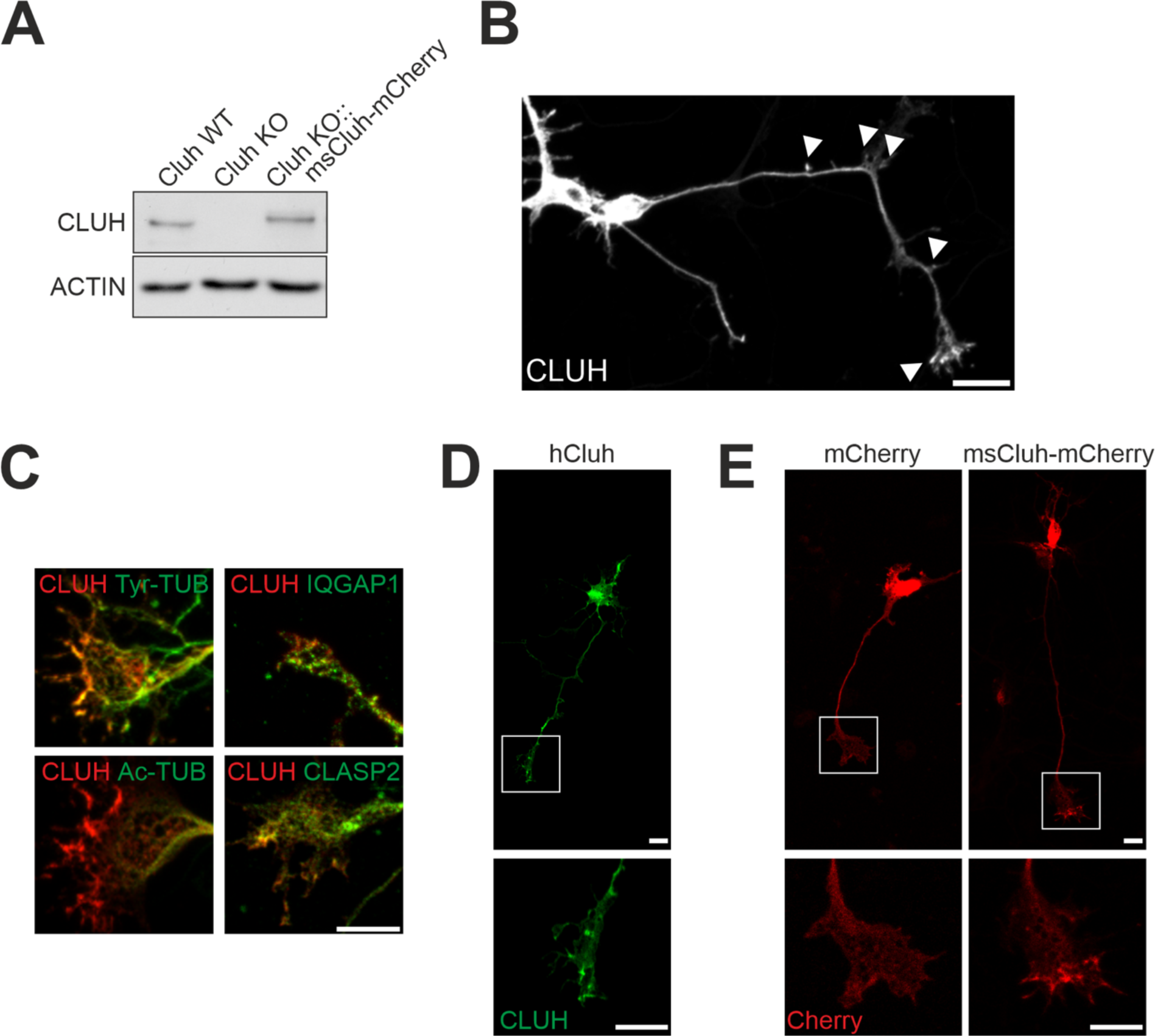
Overexpressed Cluh-mCherry localizes to somas, axons and GCs (**A**) Western blot depicting CLUH levels in WT or KO mouse embryonic fibroblasts transfected with ms*Cluh*-mCherry (ms, mouse). (**B**) Overexpression of CLUH-mCherry in a primary spinal motoneuron shows signal in the soma, in protrusions and the GCs (arrowheads). Bar, 20 µm. (**C**) GCs of primary motoneurons transfected with CLUH-mCherry (red) and stained for proteins of the cytoskeleton (green). Ac-TUB, acetylated tubulin, Tyr-TUB, tyrosinated tubulin. Bar, 10 µm. (**D**) A primary motoneuron transfected with untagged human *CLUH* and stained with an anti-human CLUH antibody (green). The boxed area containing a GC is enlarged in the lower image. Bars, 10 µm. (**E**) Primary motoneurons transfected with mCherry or ms*Cluh*-mCherry. Boxed areas showing GCs are enlarged in the lower image. Bars, 10 µm.

**Supplementary Fig. 6:**
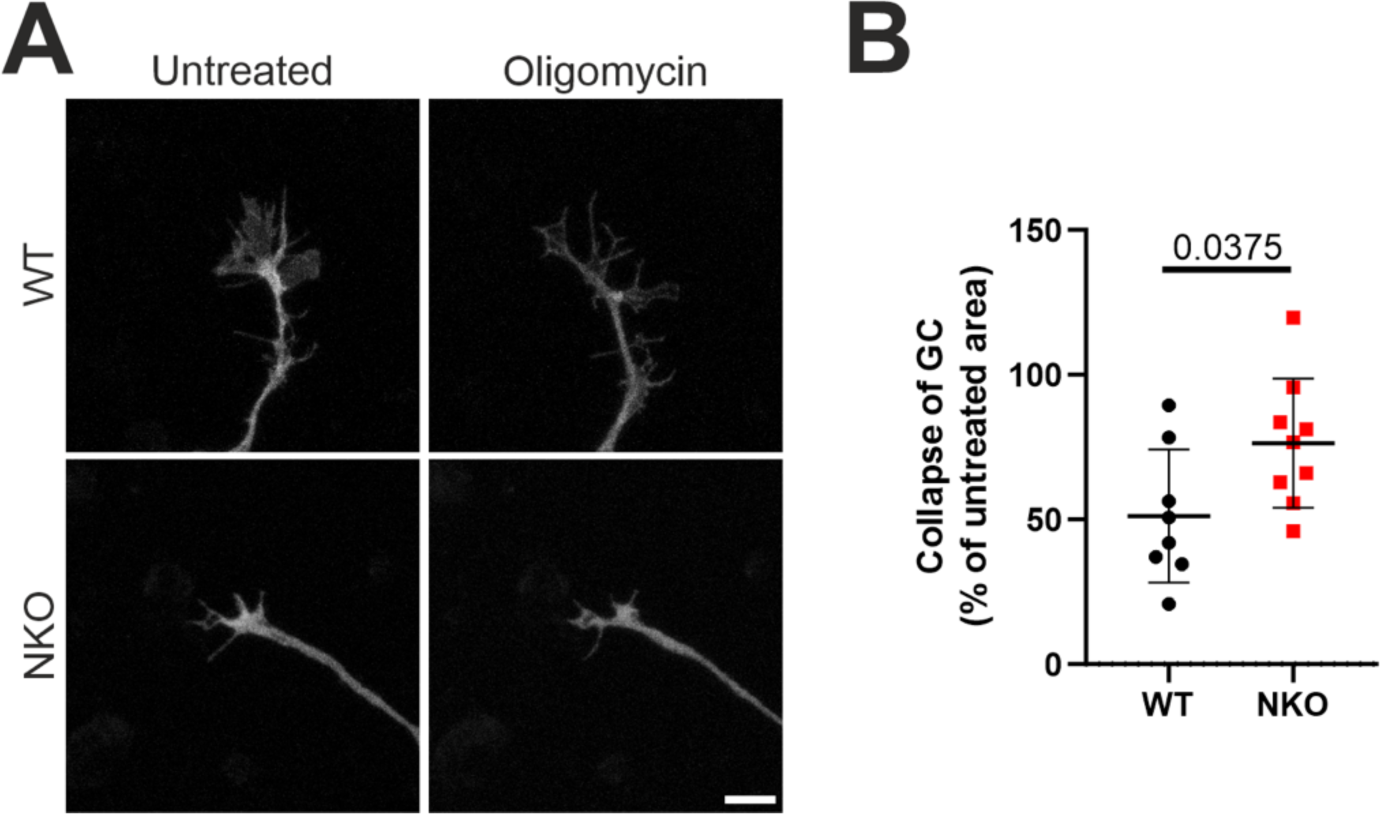
Inhibition of ATP synthase collapses WT GCs but not NKO GCs (**A**) GCs transfected with GFP (grey) were live-imaged before and after addition of oligomycin. Bar, 10 µm. (**B**) Quantification of the percentage of reduction of the area of oligomycin-treated GCs (15 min after administration) in respect to relative untreated GCs in experiments as in (A). Data represent mean ± SD of 8-9 GCs from different mice. Statistical significance was determined by Welch’s t test.

**Supplementary Fig. 7:**
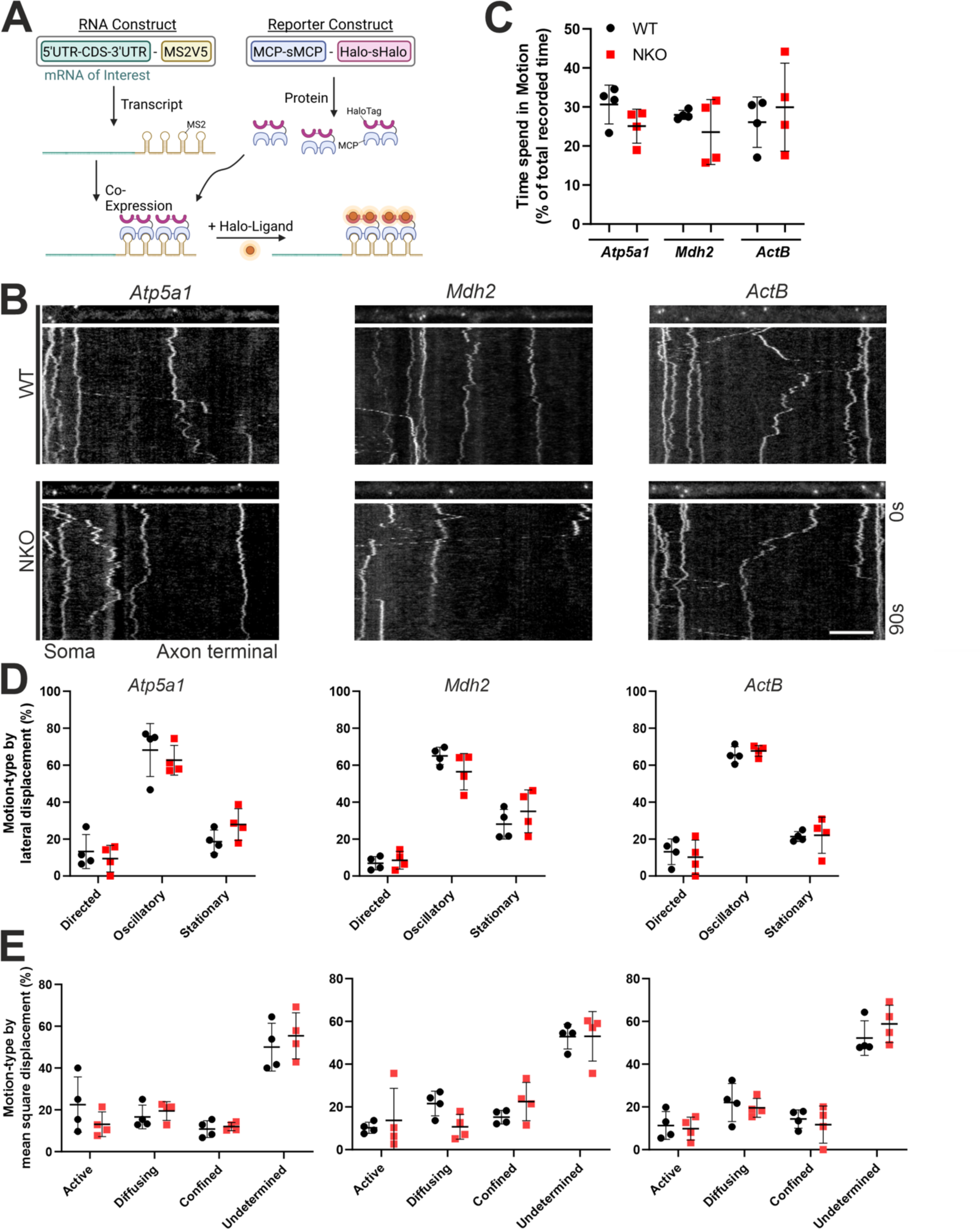
CLUH does not alter the transport of CLUH target mRNAs (**A**) The MS2-MCP system. An “RNA construct” with the target mRNA and a 3’ attached MS2V5 sequence (RNA-MS2) is co-expressed with a “reporter construct”, coding for the MCP coating protein tagged with HaloTag as the reporter protein (MCP-Halo). After the RNA construct is transcribed, MS2 stem loops form, allowing MCP-Halo to bind. Addition of Halo-Ligand enables the visualization of the mRNA. (**B**) Kymograph of *Atp5a1*, *Mdh2* and *ActB* mRNAs in axons of primary motoneurons using the MS2-MCP system depicted in (A). The first frame of the recording is shown as a straightened segment in the image above the kymograph. Bar, 5 µm. (**C**) Time spent in motion relative to the total tracked time and using a speed cut-off of more than 0.1 µm/s in experiments as in (B). (**D**) Three different motion-types (directed, oscillatory, stationary) as defined by total and maximal lateral displacement over the whole track or (**E**) by the mean square displacement in the first 25% of the track (active, diffusive, confined). Data represents the mean ± SD of 4 cultures (10-93 mRNA dots per culture).

**Supplementary Fig. 8:**
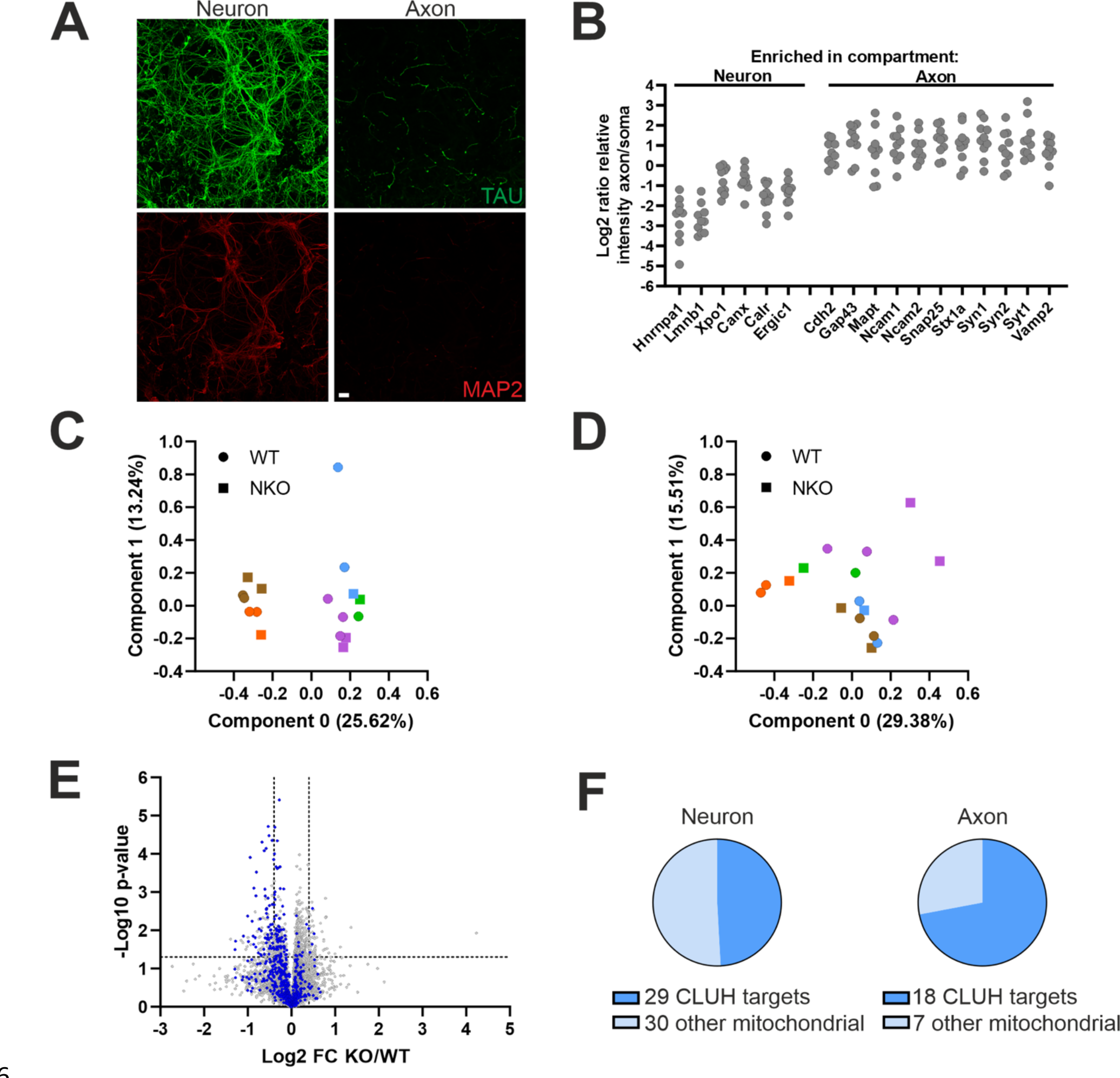
Validation of the proteomics from Boyden chambers (**A**) Neuron (left panel) and axon (right panel) compartments of Boyden chambers. Primary motoneurons are stained with MAP2 (red) and TAU (green). Bar, 25 µm. (**B**) Ratio of the protein intensities of the neuron and the axon compartments per sample. Hnrnpa1, Lmnb1 and Xpo1 were considered as proteins enriched in nuclei, Canx and Calr in ER and Ergic1 in Golgi. (**C, D**) Principal component analysis of proteomics in the neuron (C) and axon (D) compartment. Each day of collection of samples is depicted by a different colour. (**E**) Volcano plot showing protein changes in NKO versus WT neuron compartments of Boyden chambers. Mitochondrial proteins are highlighted in blue. (**F**) Pie charts depicting the percentage of mitochondrial proteins downregulated on the total of measured mitochondrial proteins which are putative CLUH targets in the neuron (left) and axon (right) compartments.

**Supplementary Fig. 9:**
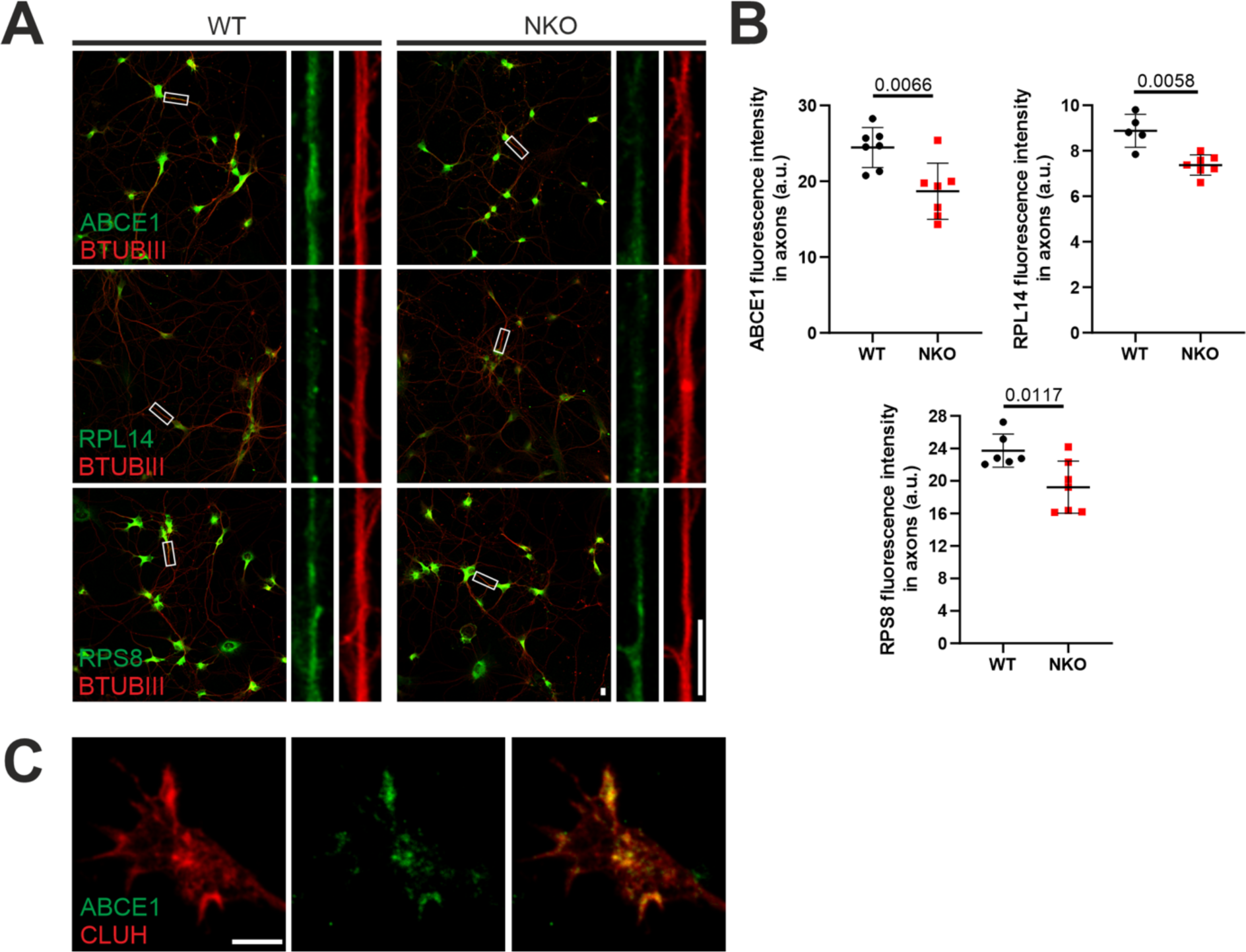
ABCE1, RPL14 and RPS8 are reduced in NKO axons (**A**) Primary spinal motoneurons stained for ABCE1, RPL14 or RPS8 (green) and β-TUBULIN III (red). Axons are straightened and magnified next to the field showing motoneurons. Bars, 10 µm. (**B**) Quantification of the intensity of the fluorescence of ABCE1 (upper left graph), RPL14 (upper right graph) and RPS8 (lower graph) in axons of experiments as in (A). Data represent the mean ± SD of 5-7 cultures (30-58 axons per culture). Statistical significance was determined by Welch’s t test. (**C**) Single confocal plane of a GC transfected with CLUH-mCherry (red) and stained for ABCE1 (green).

